# Anterior insula activity during alcohol and social reward self-administration and choice in male and female rats

**DOI:** 10.64898/2026.01.13.699207

**Authors:** Yvar van Mourik, Dustin Schetters, Ilse Bassie, Mohamad El Samadi, Huibert D. Mansvelder, Taco J. De Vries, Nathan J. Marchant

**Affiliations:** Department of Anatomy & Neurosciences, Amsterdam Neuroscience, Amsterdam University Medical Centers, Amsterdam, 1081 HZ, The Netherlands; Compulsivity Impulsivity and Attention, Amsterdam Neuroscience, Amsterdam 1081 HZ, the Netherlands; Department of Integrative Neurophysiology, Center for Neurogenomics and Cognitive Research, Amsterdam Neuroscience, Vrije Universiteit, Amsterdam 1081 HZ, the Netherlands

**Keywords:** anterior insula cortex, alcohol, social, choice, fiber photometry, addiction

## Abstract

Alcohol use disorder (AUD) is characterized by the prioritization of alcohol over healthier non-alcohol rewards, posing significant challenges for addiction treatment. Understanding the neural mechanisms driving this maladaptive preference is crucial for effective intervention. We previously showed that rats will choose alcohol over social reward in a discrete-choice task. Here, we used fiber photometry to investigate how anterior insula cortex (aIC) activity relates to choice and employed Linear Ballistic Accumulator (LBA) modelling to dissect the underlying decision processes. Male and female rats, transfected with calcium indicator jGCaMP7f in aIC, were trained to lever-press for either social reward or alcohol (20% ethanol) in alternating sessions, followed by discrete-choice sessions, and then punishment of alcohol choices. Rats developed a preference for alcohol over social reward, which was reversed when alcohol choices were punished. Model output successfully described this behaviour with the model-derived ’decision bias’ tracking preference across all phases. Photometry recordings showed that, as alcohol preference emerged, increased aIC activity during the cue period preceding alcohol choices (relative to social choices) was significantly correlated with decision bias towards alcohol. During punishment, aIC activity bias was no longer related to decision bias, despite the preference shift. These results demonstrate that aIC activity is linked to alcohol reward and choice and suggest that aIC contributes to alcohol preference by encoding a bias in the evidence accumulation process. This highlights a specific role of aIC in the cognitive mechanisms of alcohol-seeking, and its potential as a target for therapeutic interventions.

**Significance:** Understanding the neural basis of prioritizing alcohol over natural rewards is critical for combating alcohol use disorder. This study investigates how anterior insula cortex (aIC) activity in rats relates to choices between alcohol and social reward. Using fiber photometry and cognitive modelling, we found that aIC activity is not only consistently higher for alcohol-related actions but also, during the decision period prior to choice. Differential activity for alcohol versus social choices significantly correlates with a model-derived "decision bias", the speed of evidence accumulation, favouring alcohol, once preference is established. This provides novel mechanistic insight, suggesting aIC contributes to alcohol preference by encoding a bias in the decision-making process, highlighting its role in alcohol seeking and as a potential therapeutic target.

## Introduction

Alcohol use disorder (AUD) continues to impose a significant burden on individuals and society. The effects of chronic alcohol use on the body and brain are well documented (Le et al., 2001; Gilpin and Koob, 2008), but the mechanisms by which such use leads to persistent alcohol use despite negative consequences (Marchant et al., 2018; Domi et al., 2021), or to the exclusion of alternative rewards (Augier et al., 2023; Marchant et al., 2023) remains poorly understood. The role of decision making and choice in addiction is not without controversy (Hall et al., 2015; Pickard et al., 2015; Hart, 2017), in part because intact choice in addiction can be construed to argue against the brain disease model of addiction (Leshner, 1997; Heilig et al., 2021). However, disordered choice caused by the impact of chronic drug and alcohol use on the associated neurobiological systems has been proposed as a potential to addiction (Jentsch and Taylor, 1999; Goldstein and Volkow, 2011; Heatherton and Wagner, 2011; Bickel et al., 2014; Heilig et al., 2017; Heyman, 2021). Furthermore, treatment strategies incorporating choice, such as community reinforcement and contingency management, can be effective in promoting abstinence (Hunt and Azrin, 1973; Prendergast et al., 2006).

Rodent studies of choice between drug and non-drug rewards have found that rats typically show a choice preference for the non-drug reward (Lenoir et al., 2007; Cantin et al., 2010; Caprioli et al., 2015; Augier et al., 2018). Recent research has recognised the importance of social factors in addiction treatment in humans (Heilig et al., 2016). In rodents, choice for social reward can attenuate drug taking and seeking (Venniro et al., 2018). Recently, however, we found that rats choose alcohol over social reward (Marchant et al., 2023), a finding replicated in a different strain of rats (Augier et al., 2023). Furthermore, food pellets are also chosen over social reward in the discrete choice procedure (Chow et al., 2022). Thus, while for intravenously administered drugs social choice has protective value, this is not the case for orally consumed drugs like alcohol. Research directed towards understanding the neurobiology underlying choice in addiction has the potential to yield treatments which enhance behavioural allocation towards healthy, non-drug alternatives.

Lesion studies implicate the anterior insula cortex (aIC) in addiction (Naqvi et al., 2007; Joutsa et al., 2022), and fMRI studies show that exposure to alcohol-associated cues is associated with greater activation of aIC in patients (Janes et al., 2017; Janes et al., 2020). Preclinical studies have shown that aIC inactivation decreases relapse (Forget et al., 2010; Pushparaj et al., 2013; Venniro et al., 2017; Campbell et al., 2019; Joshi et al., 2020; Ghareh et al., 2022). Insula cortex activity is also related to loss of control (Rotge et al., 2017; Joshi et al., 2020) and compulsivity (Seif et al., 2013; Belin-Rauscent et al., 2016; Jones et al., 2024), and is important for decision-making during conflict (Naqvi et al., 2014; Daniel et al., 2017). Conflict in particular is a construct which is important for decision making both in choice (approach-approach conflict) and in punishment (approach-avoidance conflict) (Miller, 1971; McNally, 2021). Conflict between competing motivations can lead to ambivalence, which is a cardinal feature of addiction that has received little focus in pre-clinical research (Heather, 1998; Vandaele and Daeppen, 2022). In this study we used male and female Long-Evans rats to measure activity in aIC using fiber photometry calcium imaging, during alcohol and social reward self-administration, during choice between alcohol and social reward, and during choice between punished-alcohol and unpunished social reward. To describe the decision processes underlying choice, we used cognitive modelling (Linear Ballistic Accumulator (LBA), (Brown and Heathcote, 2008)) to quantify the latent cognitive processes guiding these choices. We estimated the speed of evidence accumulation during the different phases of the experiment, and from this derived a measure of Decision Bias to explore the relationship between aIC calcium and decision-making in choice between alcohol and social reward.

## Materials and Methods

### Subjects

We obtained 36 Long-Evans rats (18 male and 18 female), aged 8-10 weeks upon arrival (RjOrl:LE, RGD_151356971, Janvier Labs, France). All procedures were approved by the Vrije University Animal Welfare Body (Instantie voor Dierenwelzijn VU-VUmc (IvD)) and conducted under the authority of the Dutch Central Commission for Animal Research (CCD, permit #AVD11400202010449) in accordance with European law (Directive 2010/63/EU). We trained the rats in two cohorts, the first cohort had 12 experimental rats and 8 social partners, the second cohort had 12 experimental rats and 4 new social partners, however we also used the 8 social partners from the first cohort as social partners for the second cohort. Each experimental rat was exclusively trained with a single social partner, however some social partners served in this role to multiple experimental rats.

### Housing conditions

Behavioural tests were conducted during the dark phase of the rat’s diurnal cycle (12h/12h). Food and water were available ad libitum. Upon arrival, we housed the rats in groups of 4, prior to surgery, typically 2 weeks in duration. We then selected one of those rats to be the ‘Social Partner’ for the remaining 3 ‘Experimental Rats’ for cohort 1. For cohort 2, the rats were housed as 2 ‘Experimental’ and 2 ‘ Social Partner’ rats. Throughout the experiment, each experimental rat was paired with same social partner rat. Social partners were then housed together in groups of 4, and the experimental rats were single housed for the remainder of the experiment.

### Apparatus

We used Med Associates operant chambers for these experiments (See **Figure S1**). Each operant chamber consisted of a main compartment (for the experimental rat) and a smaller adjacent compartment (for the social partner), with a grid floor in both compartments. One wall of the main compartment was for social reward. There was a retractable lever, and above the lever a white cue light was the discriminative stimulus and a red cue light was used as the conditioned stimulus. There was also an automated guillotine door separating the main compartment and the adjacent compartment with a grid panel inserted so that the social partner could not enter the main compartment of the operant chamber (Venniro et al., 2019). The other wall of the main compartment was used for alcohol self-administration. This was equipped with another retractable lever, and above the lever a house light was the discriminative stimulus and a red cue light was used as the conditioned stimulus. There was also a receptacle to receive the alcohol infusions for the rat to drink out of. Alcohol was infused into the receptacle via tubing (2806 tygonslang e3603 2, 4(ID) x 4 (mm)) connected to a 20ml syringe controlled by a Razel pump. All the protocols in the chambers were ran by MedPC program and the fan remained on for the whole duration. During self-administration and choice, all operant chambers had a custom grid floor with 26 rods, however in order to connect the shock harness, we changed to the grid floor with 19 rods (VFC-005) for the punished-alcohol choice sessions. We were unable to counterbalance which side of the chamber was alcohol or social reward.

For fiber photometry, excitation and emission light was relayed to and from the animal via optical fibre patch cord (0.48 NA, 400 µm flat tip; Doric Lenses). Blue excitation light (490 or 470nm LED [M490F2 or M470F2, Thorlabs]) was modulated at 211 Hz and passed through a 460-490nm filter (Doric Lenses), while isosbestic light (405nm LED [M405F1, Thorlabs]) was modulated at 531 Hz and passed through a filter cube (Doric Lenses). GCaMP7f fluorescence was passed through a 500-550nm emission filter (Doric Lenses) and onto a photoreceiver (Newport 2151). Light intensity at the tip of the fiber was measured before training sessions 2 times a week and kept at 40-50 uW. A real-time processor (RZ5P, Tucker Davis Technologies) controlled excitation lights, demodulated fluorescence signals and received timestamps of behavioural events. Data was saved at 1017.25Hz and analyzed with Matlab scripts.

### Alcohol

We prepared 20% ethanol by diluting 70% ethanol (VWR International) with water (v/v). The first cohort of rats (12 Experimental Rats) had home-cage water that was acidified (pH = 2.7), and the operant chamber alcohol was mixed with sterile water. The second cohort of rats (12 Experimental Rats) had home-cage water that was autoclaved and thus not acidified and the operant chamber alcohol was mixed with autoclaved water.

### Viral vectors

We purchased premade viral vectors from the University of Zurich viral vector core: ssAAV-9/2-mCaMKIIα-jGCaMP7f-WPRE-bGHp(A) (**jGCaMP7f**). The titer injected was: jGCaMP7f, 5.0x10^12 gc/ml.

### Surgery

Both one day and thirty minutes prior to surgery we injected rats with the analgesic Rymadil® (5 mg/kg, s.c.; Merial, Velserbroek, The Netherlands), we also administered Buprenorphine 30 minutes prior to surgery (0.01 mg/kg, s.c.). Surgery was performed under isoflurane gas anesthesia (PCH; Haarlem). We placed the anesthetized rat in a stereotactic frame (David Kopf Instruments, Tujunga, CA) and injected Lidocaine 10 mg/kg (Fresenius Kabi, Huis ter Heide, The Netherlands; RVG 51674) into the incision site prior to the incision. A craniotomy above aIC was performed, and 0.5 µl of AAV solution was injected bilaterally into aIC (AP: +2.8, ML: +4.0 (2° angle), DV: -5.8 mm from Bregma) over 5 min. The needle was left in place for an additional 5 min. Two 400μm optic fibers (Doric Lenses) were then implanted above each aIC (AP: +2.8, ML: +3.8 (0° angle), DV: -5.5 mm from Bregma), and secured to the skull using dental cement (Antibiotic Simplex; Stryker, orthopaedics) and jewelers screws (Jeveka; M1x2 stainless steel screws). Rymadil (5 mg/kg; s.c.) was administered for 2 days after the surgery. Two rats (1M/1F) were given only unilateral aIC injection of AAV, with bilateral fiber implants, to test the extent to which contralateral projections could be contributing to the signal recorded in each hemisphere.

### Behavioural procedure

#### Phase 1: Self-administration

For both alcohol and social reward we first gave the rats a session where non-contingent rewards were delivered, and no levers were present in the chamber. For alcohol, this 60-minute session consisted of an alcohol reward (0.1mL) being delivered every 3 minutes (20 in total), and each delivery also coincided with 5 second presentation of the alcohol-associated cue (no lever press). For social, this 60-minute session consisted of the guillotine door being opened for 60 seconds, paired with the red cue-light for the first 20 seconds, every 5 minutes.

For alcohol self-administration training, we trained the rats to lever press for alcohol reward (FR1) for a total of 10 x 30-minute sessions. The alcohol-lever was inserted into the chamber, and the alcohol discriminative stimulus was turned on, throughout the whole session (note the social-lever was not inserted into the chamber during these sessions). One press on the lever resulted in 0.1ml of alcohol into the receptacle, the red cue light above this lever being turned on for 5 seconds. During this time additional lever presses resulted in no alcohol infusion (i.e. time-out period). Any alcohol remaining in the magazine after session was recorded. We calculated total alcohol consumption in grams per kilogram body weight by multiplying total alcohol rewards by 0.1 (mL), subtracting alcohol remaining, and dividing this by body weight (Kg). For social self-administration training, we also trained the rats for a total of 10 x 60-minute sessions. Before each session, we first placed the social partner rat into the adjacent social holding component of the operant chamber, and then the experimental rat (of the same sex) was placed into the main component of the operant chamber. The social-lever was inserted into the chamber, and the social discriminative stimulus was turned on, throughout the whole session (note the alcohol-lever was not inserted into the chamber during these sessions). A single lever press on the social-lever resulted in 60 seconds of social reward signalled by the opening of the guillotine door and the red cue-light above this lever turned on for 20 seconds. Responses on the social lever during this period were recorded but had no programmed consequence.

In this experiment we trained alcohol and social reward self-administration in alternating sessions over successive days (Augier et al., 2023). Alcohol self-administration was the first session, and social self-administration training followed. In the first cohort of rats, one alcohol self-administration session was excluded (session 7), because this was the first day the rats were tethered to the optic fiber patch cord, and there was a significant reduction in alcohol self-administration on this day (data not shown).

#### Phase 2: Choice tests

**Figures 1E, 1F** describe the choice session procedure. We tested choice between alcohol and social reward over 6-8 sessions after the last self-administration session. Our choice test procedure was modified slightly from our previous study (Marchant et al., 2023). Each choice test session was comprised of 20 trials, each of 4 minutes duration. Each trial started with presentation of both the alcohol and social reward discriminative cues followed 10 seconds later by insertion of both levers into the chamber. The levers were programmed to remain inserted into the chamber for two minutes, or until one was pressed. After a lever press, both levers retracted and the corresponding outcome was delivered (either 0.1ml alcohol infusion and 5 second alcohol-associated CS, or 60 seconds social interaction and 20 second social-associated CS). Because the choice session is divided into 20 x 4 minute blocks, the ITI is variable depending on the response latency of the choice, but it does not vary depending on the outcome (alcohol or social). If no lever was pressed after two minutes the discriminative cues were turned off and both levers were retracted. Thus, if the rat pressed either lever at the beginning of the trial, then the ITI would be 3 minutes 50 seconds, but if they did not press at all (omission) then the cues are turned off and levers retracted for 1 minute and 50 seconds.

**Figure 1.**
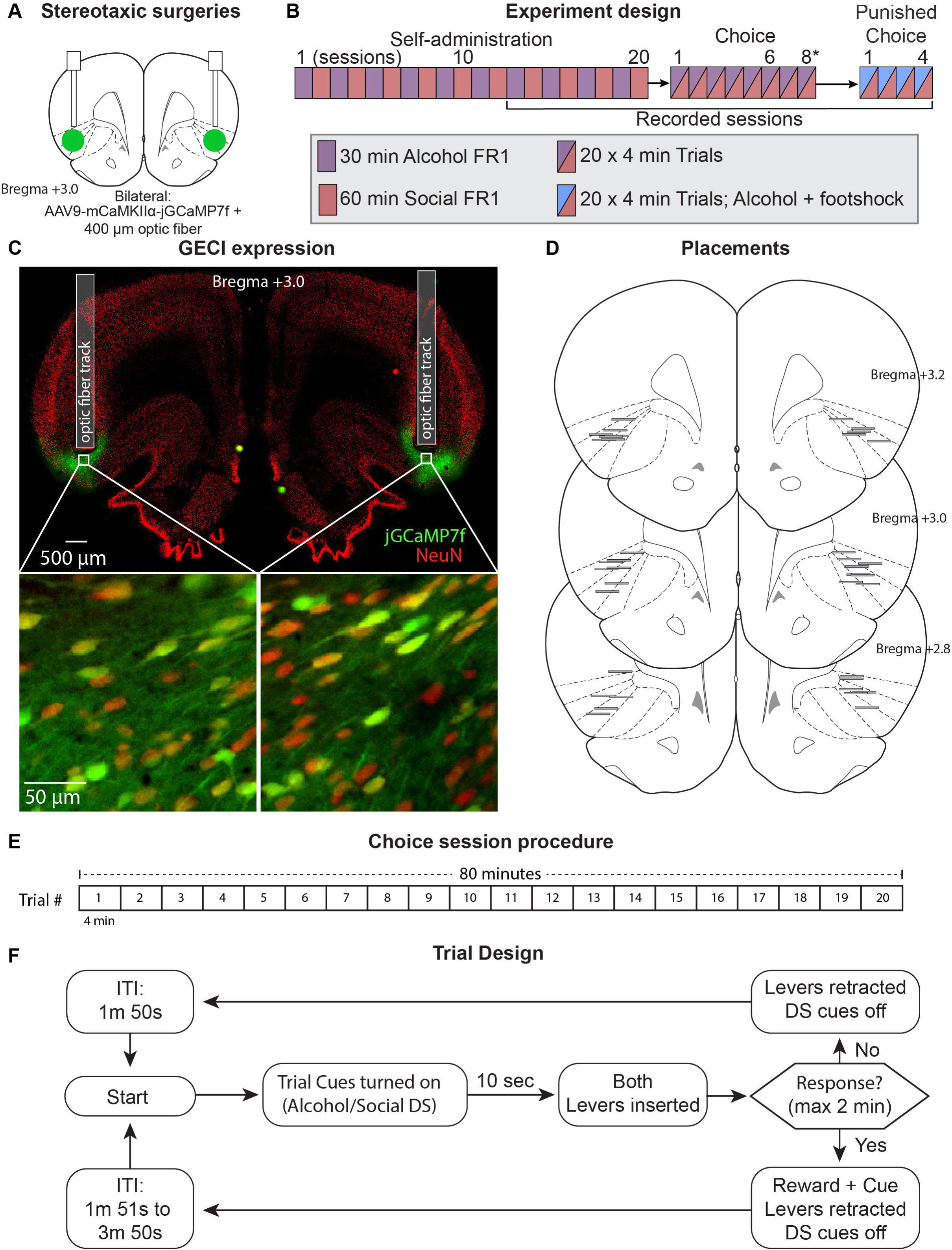
Experimental outline. (**A**) Outline viral targeting strategy. We injected AAV encoding jGCaMP7f into aIC, and implanted fiber optics above. (**B**) Experiment Design. We trained rats in alternating sessions of alcohol or social reward. During choice, the first cohort (n=12) received 6 choice sessions, the second cohort (n=11) received 8 choice sessions. All rats received 4 punished-alcohol choice sessions, where alcohol lever press resulted in 0.3 mA intensity, 0.5 sec duration shock. (**C**) Representative jGCaMP7f and fiber placement in aIC. (**D**) Depiction of fiber placement for all rats, horizontal lines represent the base of the fiber for rats included in the analysis. (**E**) Depiction of the choice session procedure. Each choice session consisted of 20 x 4-minute trials. (**F**) Trial Design. A trial begins with onset of the discriminative stimuli (DS) for 10 seconds and then both levers are inserted. After a response, both levers are retracted, both DS are turned off, and the relevant reward is delivered. If no response is made in 2 minutes, then both levers are retracted, both DS are turned off. *ITI, inter-trial-interval*; *DS, Discriminative Stimulus*.

#### Phase 3: Punished-alcohol choice tests

The timing parameters of this phase are identical to the previous choice phase. When an alcohol choice was made, the outcome was 0.1ml alcohol infusion into the magazine, 5 second presentation of the alcohol-associated CS, and an electric foot shock of 0.5 second duration and 0.3 mA intensity. Social choice resulted in the same outcome as before, with no foot shock. We gave all rats 4 sessions in this phase, recording left and right hemisphere each twice. We chose the intensity of 0.3 mA initially because we observe substantial individual variability in single-lever operant conditions (e.g. (McDonald et al., 2024)). We were surprised to find that this intensity was sufficient to fully shift preference to social reward for both male and female (except 2 rats) by the fourth session, and as such used this intensity for the second cohort of rats.

### Photometry recording sessions

We recorded either left or right hemisphere aIC for all rats over alternating sessions of alcohol and social self-administration, choice, and punished-alcohol choice. The self-administration sessions were only recorded for the last 8 sessions, resulting in 4 sessions each of alcohol or social recorded (2 left and 2 right for each). All choice and punished-alcohol choice sessions were recorded.

### jGCaMP7f expression and fiber placement validation

We deeply anesthetized rats with isoflurane and Euthasol® injection (i.p.) and transcardially perfused them with ∼100 ml of normal saline followed by ∼400 ml of 4% paraformaldehyde in 0.1M sodium phosphate (pH 7.4). The brains were removed and post-fixed for 2 h, and then submerged in 30% sucrose in 0.1M PBS for 48 h at 4°C. Brains were then frozen on dry ice, and coronal sections were cut (40 µm) using a Leica Microsystems cryostat and stored in 30% sucrose in 1.0M PBS stored at -20°C.

Immunohistochemical procedures are based on our previously published work (Marchant et al., 2009; Marchant et al., 2010; Campbell et al., 2019; Ghareh et al., 2022). We selected a 1-in-4 series and first rinsed free-floating sections (3 x 10 minutes) before incubation in PBS containing 0.5% Triton-X and 10% Normal Donkey Serum (NDS) and incubated for at least 48 h at 4°C in mouse anti-NeuN primary antibody (1:1000; Chemicon, MAB377). Sections were then repeatedly washed with PBS and incubated for 2-4 h in PBS + 0.5% Triton-X with 2% NDS and donkey anti-mouse secondary antibody DyLight 649 (1:500; Jackson ImmunoResearch, 715-495-150). After another series of washes in PBS, slices were stained with DAPI (0.1 ug/ml) for 10 min prior to mounting onto gelatin-coated glass slides, air-drying and cover-slipping with Mowiol and DABCO.

Slides were all imaged on a VectraPolaris slide scanner (VUmc imaging core) at 10x magnification, and QuPath was used for image analysis (Bankhead et al., 2017). Images containing aIC, from Bregma + 4.2 mm to +2.5 mm were identified and the boundary of expression for each rat was plotted onto the respective Paxinos and Watson atlas (Paxinos and Watson, 2008).

### Behavioral Data Analysis: Linear Ballistic Accumulator (LBA) Modelling

To investigate the latent decision processes underlying choice behaviour, we fitted reaction time (RT) and choice data from each rat and experimental session (’early’, ’late’, ’punishment’) to a hierarchical Linear Ballistic Accumulator (LBA) model (Brown and Heathcote, 2008). The LBA is a well-established sequential sampling model that describes decision-making as a race between independent accumulators, one for each response option (in this study, alcohol and social reward).

The first accumulator to reach a decision threshold determines the choice made and the decision time for that trial. By implementing the model in a hierarchical framework, individual subject- and session-level parameters are estimated as being drawn from overarching group-level distributions. This approach allows the model to borrow strength across subjects and sessions, separating true variability from estimation noise and resulting in more stable and reliable parameter estimates.

### LBA Model Parameters

The LBA model characterizes choice and RT through several key parameters, which were estimated for each rat and session: **Mean Accumulation Rate (v)**: The average rate at which evidence accumulates for a specific response option (i.e., v_alcohol_, v_social_). For each trial, the actual accumulation rate for an accumulator is sampled from a normal distribution with mean (v_option_) and standard deviation (s). Higher mean accumulation rates lead to faster and more frequent choices of that option.

### Between-Trial Variability in Accumulation Rate (s)

This parameter captures trial-to-trial variability in the speed of evidence accumulation. Following common practice for model identifiability and comparison with previous work (Annis et al., 2017; Choi et al., 2022), we fixed s to 1. This allows other parameters, particularly the mean accumulation rates, to scale accordingly and represent the signal-to-noise ratio of the accumulation process. **Start-Point Variability (A)**: The upper bound of a uniform distribution (U[0,A]) from which the starting point of evidence accumulation for each accumulator is randomly sampled on each trial. Larger A indicates more variability in the initial state of evidence. **Decision Threshold (b)**: The amount of evidence required for an accumulator to trigger a response. In our parameterization, b was derived from two estimated parameters: a relative threshold component (k) and the start-point variability (A), such that b=k+A. This formulation ensures that the decision threshold b is always greater than any possible starting point (k is constrained to be positive). **Non-Decision Time (tau)**: This parameter accounts for time consumed by perceptual and motor processes that are peripheral to the evidence accumulation and decision stage (e.g., stimulus encoding, response execution). The total observed RT on a given trial is the sum of the decision time (determined by the race) and tau.

### Hierarchical Model Specification and Comparison

To systematically determine which cognitive mechanisms rats used to adapt their behaviour, we employed a ’power set’ approach. We specified three core mechanisms that could vary across the experimental sessions: 1) Stimulus Bias (S), a change in the relative evidence accumulation rates (v_alcohol vs. v_social); 2) Caution (C), a change in the decision threshold, parameterized via start-point variability (A); and 3) Response Bias (R), an asymmetry in the starting point of accumulation, parameterized via the relative threshold (k). We then constructed and fit eight hierarchical LBA models, representing all possible combinations of these mechanisms (i.e., a baseline model with no mechanisms varying, three single-mechanism models, three double-mechanism models, and one triple-mechanism model)

We performed a formal model comparison using Leave-One-Out Cross-Validation (LOO-CV), a robust method for estimating out-of-sample predictive accuracy. The results, assessed via the expected log predictive density (ELPD), indicated that model M_SC, in which Stimulus Bias (S) and Caution (C) were allowed to vary across sessions, provided the best and most parsimonious account of the data (see Supplementary Materials). Therefore, all subsequent analyses and interpretations are based on the posterior parameter estimates from this winning model.

### Parameter Priors for the Winning Hierarchical Model (M_SC)

We fitted the LBA model within a Bayesian framework using PyMC (Salvatier et al., 2016). Priors for the group-level model parameters were chosen to be weakly informative: <u>Group-level Means (μ):</u> For parameters that varied across sessions in the winning M_SC model (v_alcohol, v_social, and A), separate group-level means were estimated for each session. Parameters held constant across sessions for each subject (k and τ) were estimated with a single group-level mean. <u>Group-level Standard Deviations (σ):</u> HalfNormal distributions were used for the standard deviations of all group-level parameters. <u>Subject/Session-level Offsets:</u> Individual subject- and session-level parameters were modelled as offsets from the group-level mean, drawn from a Normal distribution with a standard deviation determined by the corresponding group-level σ. This non-centered parameterization improves sampling efficiency. <u>Non-Decision Time (τ):</u> For each subject, the τ parameter was bounded to be less than the minimum observed RT for that subject, ensuring plausibility.

### Derived Parameters

From the primary posterior distributions of the winning model, we derived two key cognitive metrics: *<u>Response Caution:</u>* Defined as *b* − *A*/2, reflecting the average distance from the mean starting point to the threshold. *<u>Decision Bias:</u>* We define this metric as the accumulation rate difference (v_alcohol_ and v_social_). This value reflects the relative speed and efficiency of evidence accumulation toward one choice over the other, rather than a pre-decisional bias in the starting point.

### Model Fitting and Convergence

The hierarchical LBA model was fitted to the complete dataset of all trials from all rats. We used the No-U-Turn Sampler (NUTS;) (Hoffman and Gelman, 2014), as implemented in PyMC. We ran 4 parallel chains, each with 1500 tuning (warm-up) steps and 2000 sampling steps, resulting in 8000 posterior samples per parameter. Convergence of the MCMC chains was assessed using the R-hat (R^) statistic, with R^ < 1.01 indicating successful convergence (Gelman and Rubin, 1992), and the effective sample size (ESS), with ESS > 400 considered adequate (Gelman et al., 2013). Trace plots were also visually inspected for chain mixing and stationarity.

<u>Model Evaluation:</u> To evaluate how well the fitted models could replicate observed behavioural patterns, posterior predictive checks (PPCs) were performed for each converged model. This involved simulating 500 new datasets, each with the same number of trials as the original rat-session dataset, by drawing 500 parameter sets from the joint posterior distribution of the fitted model. We then compared the distributions of these simulated reaction times (separated by choice option) and the simulated choice proportions against the actual observed data. All analyses were performed using Python (version 3.10.15) and the PyMC (version 5.18.2), ArviZ (version 0.20.0), pandas (2.2.3), and NumPy (1.26.4) libraries. The scripts are available at: https://github.com/njmarchant/LBA-Alcohol_Social_Choice

### Statistics

<u>Behaviour:</u> All data was analyzed using IBM SPSS V21. Phases were analyzed separately, and in all tests, we compared male and female rats using Sex as a between-subjects factor. During self-administration the dependent variables were the total number of active lever presses, and total number of alcohol reward deliveries or social reward opportunities (guillotine door open for 60 seconds). For the choice tests, dependent variables were the total choice of either alcohol or social reward. To analyse we used repeated measures analysis of variance (ANOVA), using the within-session factor Session where appropriate.

Preference score was calculated using the formula: (Alcohol – Social)/(Alcohol + Social) resulting in values that range from +1 (full **<u>alcohol</u>** preference) to –1 (full **<u>social</u>** preference). To calculate the preference scores for ‘early’ and ‘late’ choice sessions, we averaged the preference score of the two sessions for each rat. For the punished-alcohol choice sessions we averaged the preference score of all four sessions. All comparisons were made using repeated-measures t-tests.

Latency data was not normally distributed, and therefore we used Kruskal–Wallis one-way analysis of variance to compare latencies for alcohol or social choice in the different sessions in a single analysis. We report the relevant post-hoc test output both uncorrected and Bonferroni corrected for multiple tests. For ’late choice’ latencies we used the last two choice sessions of phase 2, thus for cohort 1 this was sessions 5 and 6, and for cohort 2 this was sessions 7 and 8. To calculate mean latencies, omitted trials were simply excluded.

<u>Photometry:</u> Recorded signals were first downsampled by a factor of 64, giving a final sampling rate of 15.89 Hz. The 405nm isosbestic signal was fit to the 470/490nm calcium-dependent signal using a first order polynomial regression. A normalized, motion-artefact-corrected ΔF/F was then calculated as follows: ΔF/F = (490nm signal − fitted 405nm signal)/fitted 405nm signal. The resulting ΔF/F was then detrended via a 90s moving average, and low-pass filtered at 3Hz. Several different epochs were selected for analysis. In self-administration, ΔF/F from 5s before lever press to 10s after were collated. To avoid duplicate traces due to overlapping epochs in self-administration, we only included the reinforced active lever presses, and thus all time-out responses were not analysed. These traces were then baseline-corrected using the baseline time-period -5 seconds to -3 seconds prior to the reinforced lever press. We converted the data into z-scores by subtracting the mean baseline activity and dividing by the standard deviation. In choice, epochs around the lever press (choice) and around the trial start were separately analysed. For analysis based on the choice response, ΔF/F from 20s before the lever press to 20s after were collated. For analysis based on the trial start, ΔF/F from 10s before the cues turning on to 30s after were collated. These traces were then baseline-corrected and converted into z-scores with the same method described above, using the time-period of -10 seconds to -5 seconds before trial as the baseline for analyses centred on trial start, and -20 seconds to -15 seconds before the response for analyses centred on the response.

In self-administration, traces from the different sessions (alcohol or social) were compared. For analysis of the choice data, we compared choice outcome (response) and decision period (trial start with cues) separately. For the choice we grouped traces by choice outcome: alcohol, social, or omission. For the trial start we grouped traces around the response that was eventually made (alcohol, social, or omission). We used two approaches to analyse the resulting traces: bootstrapped confidence intervals and permutation tests. The rationale for each is described in detail here: (Jean-Richard-Dit-Bressel et al., 2020). The output of every statistical test we conducted is available in the raw data files.

Bootstrapped confidence intervals were used to determine whether calcium activity was significantly different from baseline (ΔF/F = 0). A distribution of bootstrapped means was obtained by randomly sampling from traces with replacement (*n* traces for that response type; 5000 iterations). A 95% confidence interval was obtained from the 2.5^th^ and 97.5^th^ percentiles of the bootstrap distribution, which was then expanded by a factor of sqrt(n/(n – 1)) to account for narrowness bias.

Permutation tests were used to compare calcium activity between the different groupings. Observed differences were compared against a distribution of 1000 random permutations (difference between randomly regrouped traces) to obtain a p-value per time point. Alpha of 0.05 was Bonferroni-corrected based on the number of comparison conditions, resulting in alpha of 0.01 for comparisons between 3 conditions. For both bootstrap and permutation tests, only periods that were continuously significant for at least 0.5s were identified as significant.

The output of these tests are displayed in the figures below the traces, where the presence of a significance bar indicates significance during that time period. Not all statistical tests are reported in the figures, however we have reported the time windows where the statistical tests yielded significant outputs in for every test in Tables 1-3. The scripts are available at: https://github.com/njmarchant/Photom-Alcohol_Social_Choice

## Results

Figure 1 shows the experimental outline and targeting strategies for calcium imaging with fiber photometry in aIC. We targeted the anterior insula cortex bilaterally with AAV encoding jGCaMPf driven under the CaMKIIa promoter (Figure 1A), Figure 1C shows example expression and Figure 1D shows the fiber placements verified after the experiment. Figure 1B shows the experimental design, we used an alternating training schedule (Augier et al., 2023), where rats were trained in separate days to self-administer on an FR1 schedule either alcohol (20% ethanol in water, 3 second time-out) for 30 minutes or social reward (60 second open door separating the experimental rat and social partner) for 60 minutes. Recording sessions were made on the final 4 self-administration sessions, all choice sessions, and all punishment sessions.

### Alcohol and social reward self-administration

Figure 2 shows the self-administration data where only one lever was available for the whole session. Figure 2A shows the total number of rewards that were delivered in each session, and Figure 2B shows the total number of lever presses. For alcohol self-administration, we found a main effect of Session for rewards (F(9,198)=5.057; p<0.001) and for lever presses (F(9,198)=4.180; p<0.001).

**Figure 2.**
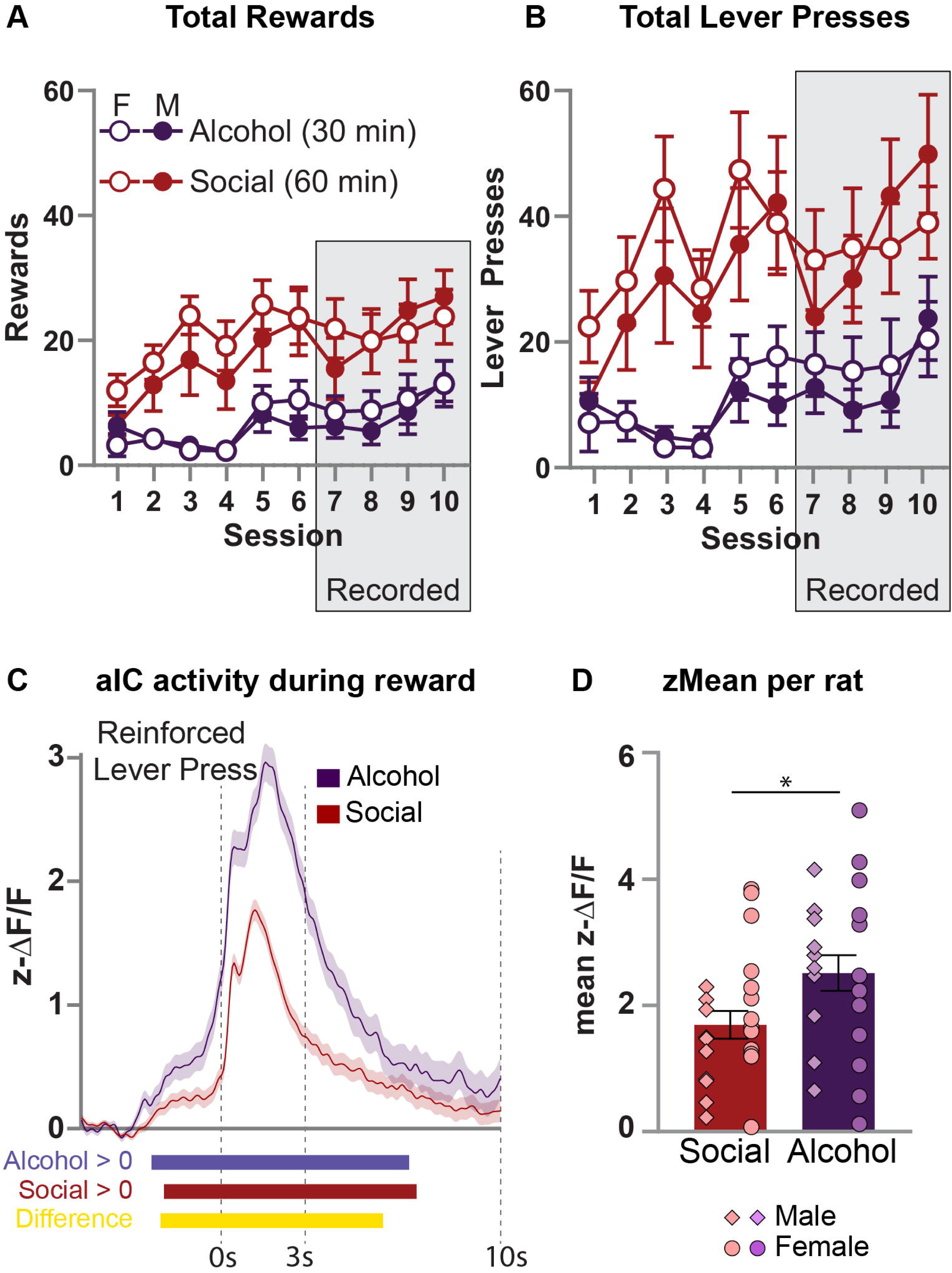
Activity in aIC during alcohol and social reward self-administration sessions. (**A**) Total Rewards, and (**B**) Total Lever Presses, during the alcohol or social reward self-administration sessions. (**C**) Traces depicting aIC activity surrounding the reinforced lever presses (0s) during the self-administration sessions. Only reinforced lever presses are represented here. (**D**) Mean (±SEM) of z-Scored activity during the 3 second outcome period during self-administration. Table 1 shows the time-windows where the statistical tests were significant. *, p<0.05. n=11 males; diamond shapes, n=12 females, circle shapes.

There was no effect of Sex in these measures (F(1,22)<1; p>0.05). Analysis of alcohol consumed (g/Kg; data not shown) revealed a main effect of Session (F(1,22)=5.25; p<0.001) but no effect of Sex (F(1,22)<1; p>0.05). Alcohol consumed (g/Kg) in the final session was Male: 0.54(±0.45) and Female: 0.83(±0.89).

For social reward self-administration, we found a main effect of Session for rewards (F(9,198)=7.139; p<0.001) and for lever presses (F(9,198)=5.317; p<0.001). There was no effect of Sex in these measures (F(1,22)<1; p>0.05). To compare alcohol and social reward, we converted the data to the rate of responding per minute and found an effect of Reward Type on Total Rewards (F(1,23)=4.377; p=0.048) but not on Total Lever Presses (F(1,23)=4.128; p=0.054). Comparable analysis on the last 4 sessions of each (the recorded sessions) found no effect of Reward Type on Total Rewards (F(1,23)<1; p>0.05) or Total Lever Presses (F(1,23)<1; p>0.05).

**Figure 2C** shows the traces depicting calcium transients recorded in aIC centred around the reinforced lever presses. Bootstrapping analysis revealed that aIC activity was higher than baseline both prior to and after the lever press. Comparison between aIC activity for alcohol and social reward using permutation tests revealed a significant difference. Summary statistics using the mean of the z-scored recorded calcium during the 3 seconds after a reinforced lever press are shown in **Figure 2D**. There is a significant difference between Alcohol and Social (Alcohol mean 2.43 ± 1.31, Social mean 1.66 ± 1.04; paired t-test Alcohol vs. Social: t(21)=-2.56, p=0.019). These data show that while aIC activity is increased relative to baseline for both alcohol and social rewards, the magnitude of activation is higher for alcohol.

### Discrete choice between alcohol and social reward

**Figure 1E** shows the choice session procedure. We used a comparable procedure to our previous work (Marchant et al., 2023), however rather than 15 x 8-minute trials we gave the rats 20 x 4-minute trials. Figure 3A shows the choice data from the 8 sessions (cohort 1 (n=12) received 6 sessions, cohort 2 (n=11) received 8 sessions). One male rat (from cohort 2) was removed from the dataset because he made only 2 choice responses in 8 choice sessions. We found that over the course of the training a choice preference for alcohol emerged. Analysis of the choice responses revealed a significant Choice x Session interaction (F(5,105)=5.065; p<0.001) including both cohorts from c1 to c6. Analysis of cohort 2 from c1 to c8 also revealed a significant Choice x Session interaction (F(7,63)=10.602; p<0.001). **Figure 3B** shows preference score for the first two choice sessions (Early: c1-c2 for both cohorts), and the last two choice sessions (Late: c5-c6 for cohort 1, c7-c8 for cohort 2). Paired t-test revealed a significant difference (t(22)=3.437; p=0.0024) reflecting the emergence of choice preference for alcohol. We also analysed the preference score data with Sex as a between-subjects factor and found no Session x Sex interaction (F(1,21)=2.053; p>0.05) and no main effect of Sex (F(1,21)<1; p>0.05). One way ANOVA on alcohol consumed (g/Kg; data not shown) during the Early sessions revealed no effect of Sex (Male: 0.39±0.23; Female: 0.42±0.32; F(1,21)<1; p>0.05), however there was an effect of Sex in the Late sessions (Male: 0.55±0.27; Female: 0.91±0.0.39; F(1,21)=6.3; p=0.02).

**Figure 3.**
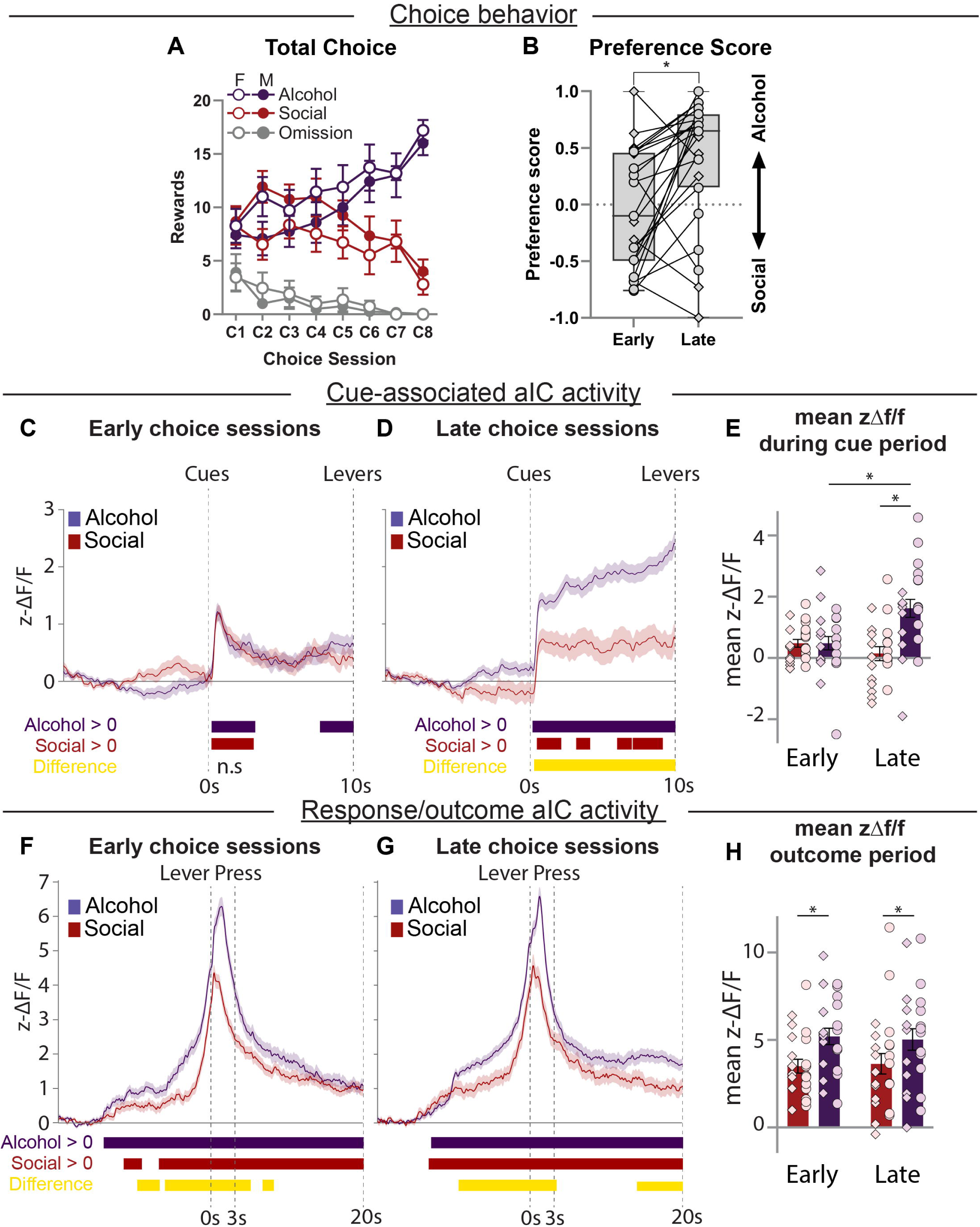
Activity in aIC during choice sessions. (**A**) Group data showing alcohol or social choices, and omissions, from the choice sessions. (**B**) Preference score comparing Early and Late choice sessions. (**C, D**) Traces depicting aIC activity centred around the trial start (0s) during all Early or Late choice trials. (**E**) Mean (±SEM) of z-Scored activity during the 10 second cue period during the Early (left) or Late (right) choice sessions. (**F, G**) Traces depicting aIC activity surrounding choice response (0s). (**H**) Mean (±SEM) of z-Scored activity during the 3 second outcome period during the Early (left) or Late (right) choice sessions. Table 2 shows the time-windows where the statistical tests were significant. *, p<0.05. n=11 males; diamond shapes, n=12 females, circle shapes.

**Figures 3C and 3D** show aIC activity centred around trial start during the Early (3C) and Late (3D) choice sessions, with the data separated into whether the trials are those where alcohol or social is chosen. During the Early choice sessions (3C), aIC activity increased relative to baseline for both choice types but was not different from each other (permutation tests). In contrast, during the Late choice sessions (3D), we found that aIC activity during the cue period is significantly higher when alcohol is chosen compared to when social is chosen. This pattern of activity is reflected in the summary statistics, shown in Fig. 3E. Within-subjects ANOVA using the factors Session (Early, Late) and Choice (Alcohol, Social) revealed a main effect of Choice (F(1,19) = 7.9, p=0.011) and a Choice by Session interaction (F(1,19)=9.20, p=0.007). Post-hoc t-tests show that aIC activity during the cue period for alcohol choices significantly increased from Early to Late choice (paired t-test Early Alcohol vs. Late Alcohol: t(21)=-3.21, p=0.004), and in the Late sessions alcohol is significantly higher than social (paired t-test Late Alcohol vs. Late Social: t(20)=3.79, p=0.001).

We performed a secondary analysis on aIC activity centred around trial start for the intermediate choice sessions (i.e. ’middle’ choice sessions are C3 and C4 sessions). These data show that there is a significant increase in aIC activity during the cue period for alcohol choices (Fig. S2B), but not social choices (Fig. S2C), in the middle choice sessions compared to early choice sessions. This is potentially interesting because there are no group differences in preference scores between these sessions (Fig. S2A; paired t-test Early vs. Middle: t(22)=0.15, p>0.05). Comparisons of aIC activity from the middle to late choice sessions show a further increase in aIC calcium towards the end of the cue period (Fig. S2E), and again no change in activity when social is chosen (Fig. S2F). Behaviourally, there was a significant change in preference score (Fig. S2D; paired t-test Middle vs. Late: t(22)=4.8, p<0.001).

**Figures 3F and 3G** show aIC activity centred around the choice response (lever press) during the Early (3F) and Late (3G) choice cessions, with the data separated into whether the choices made were alcohol or social reward. Recorded traces of aIC activity reveal comparable patterns of activity between the early and late choice sessions. In both session types, aIC activity was higher for alcohol choices leading up to the lever press, and during the outcome period alcohol was higher than social. Figure 3H shows the mean aIC activity during the 3 second period after the choice was made. We found that in both early and late choice sessions, aIC activity was higher for alcohol than social reward, and that this did not change between the sessions. Within-subjects ANOVA using the factors

Session (Early, Late) and Choice (Alcohol, Social) revealed a main effect of Choice (F(1,19) = 20.2, p<0.001) but no Choice by Session interaction (F(1,19)<1, p>0.05). Post-hoc t-tests show that aIC activity during the outcome period for alcohol is significantly higher than social in both Early and Late choice (paired t-test Early Alcohol vs. Early Social: t(21)=4.34, p<0.001; paired t-test Late Alcohol vs. Late Social: t(20)=2.17, p=0.04), and no differences between Early and Late sessions for Alcohol (Early Alcohol mean 5.27 ± 2.25, Late Alcohol mean 5.02 ± 2.88; paired t-test Early vs. Late: t(21)<1, p>0.05) or Social (Early Social mean 3.58 ± 1.84, Late Social mean 3.68 ± 2.82; paired t-test Early vs. Late: t(21)<1, p>0.05).

### Discrete choice between punished alcohol and social reward

**Figure 4A** shows the total choices made for alcohol or social during the late choice sessions and the punished-alcohol choice sessions. We found that punishment of the alcohol-reinforced choice caused a preference switch to social reward. Analysis of the choice responses comparing response from the ‘late choice’ sessions with the punished-alcohol choice sessions revealed a significant Choice x Session interaction (F(1,21)=7.243; p=0.014). One way ANOVA on alcohol consumed (g/Kg; data not shown) during the punished-alcohol choice sessions revealed no effect of Sex (Male: 0.08±0.15; Female: 0.07±0.03; F(1,21)<1; p>0.05). **Figure 4B** shows the preference score for the late choice sessions and the punished-alcohol choice sessions. Paired t-test revealed a significant difference (t(22)=-9.757; p<0.001) reflecting the emergence of choice preference for social reward.

**Figure 4.**
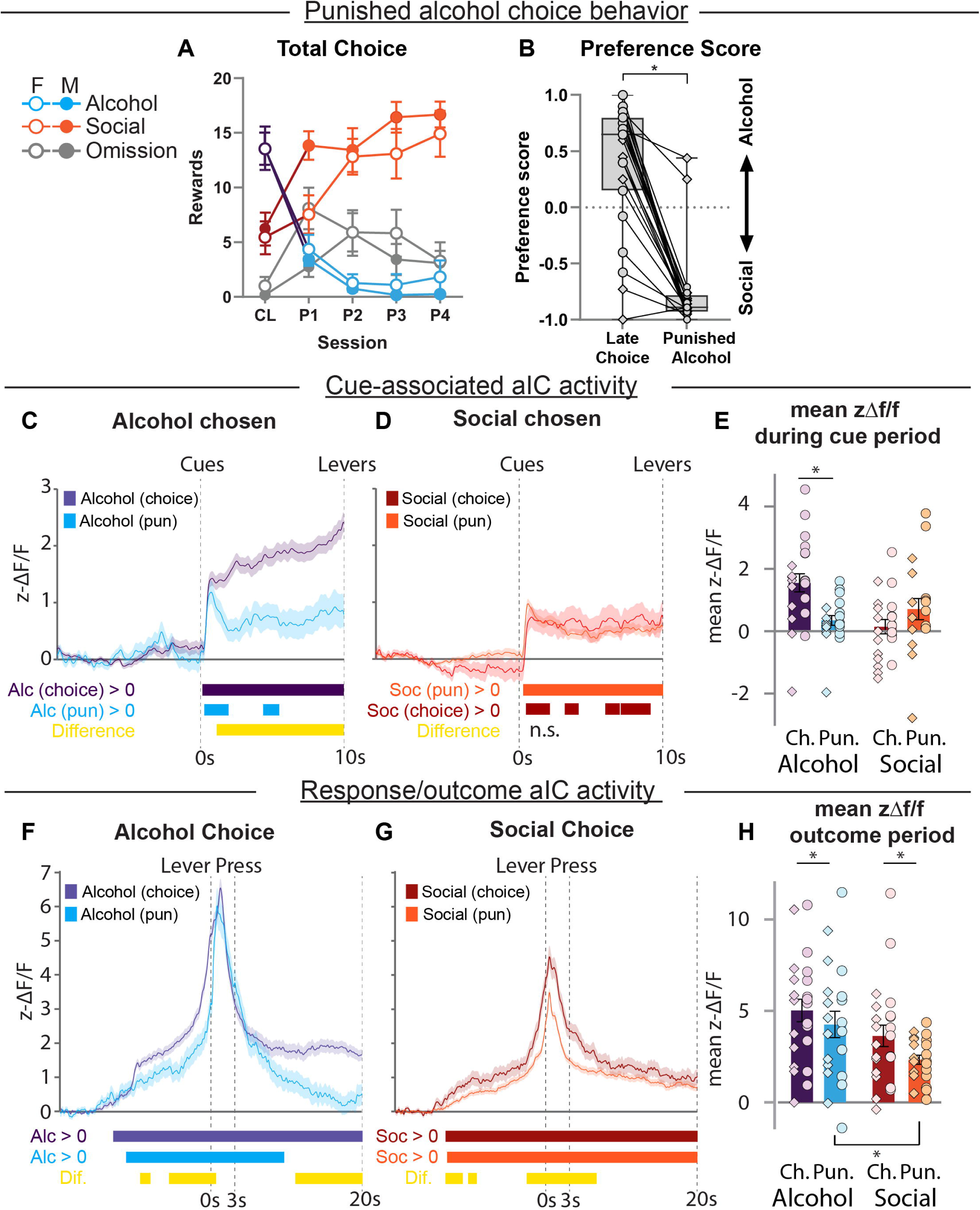
Activity in *aIC during punished-alcohol choice sessions*. (**A**) Group data showing alcohol or social choices, and omissions, from the punished-alcohol choice sessions. (**B**) Preference score comparing Late choice and Punished alcohol choice sessions. (**C, D**) Traces depicting aIC activity centred around the trial start (0s) during all punished-alcohol choice trials. (**E**) Mean (±SEM) of z-Scored activity during the 10 second cue period in trials where alcohol is chosen (left) or social is chosen (right). (**F, G**) Traces depicting aIC activity surrounding choice response (0s). (**H**) Mean (±SEM) of z-Scored activity during the 3 second outcome period in trials where alcohol is chosen (left) or social is chosen (right). Ch, Late Choice; Pun, Punished alcohol choice; Dif, Difference; Alc, Alcohol; Soc, Social. Table 3 shows the time-windows where the statistical tests were significant. *, p<0.05. n=11 males; diamond shapes, n=12 females, circle shapes.

**Figure 4C** shows aIC activity centred around trial start for trials where alcohol is chosen comparing Late choice to Punished alcohol choice sessions. **Figure 4D** shows the same comparison for trials where social was chosen. This analysis shows that aIC activity during the cue period significantly decreases in the punished alcohol choice sessions. In contrast, there is no difference in aIC activity to trials where social is chosen between the two session types. This pattern of activity is reflected in analysis of the mean aIC activity during the cue period shown in **Figure 4E**. Within-subjects ANOVA using the factors Session (Late, Pun) and Choice (Alcohol, Social) revealed a main effect of Choice (F(1,19)=9.5, p=0.006) and Session (F(1,19)=8.3, p=0.01). As well as a Choice by Session interaction (F(1,19)=7.38, p=0.014). Post-hoc t-tests show that aIC activity during the cue period for alcohol choices significantly decreased from Late choice to Punished alcohol choice (paired t-test Late Alcohol vs. Punished Alcohol: t(19)=3.48, p=0.003), and in the Late sessions there is no significant difference between (punished) alcohol and social choice trials (paired t-test Punished Alcohol vs. Social (Punish sessions): t(20)<1, p>0.05).

**Figures 4F and 4G** show aIC activity centred around the choice response during the Late (4F) and Punished alcohol (4G) choice cessions, with the data separated into whether the choices made were alcohol (+shock) or social reward. Recorded traces of aIC activity show that aIC activity leading up to alcohol choices is decreased during the punished sessions compared to the prior choice sessions, but permutation tests revealed no significant difference during the outcome period. For social choices, we found that aIC activity is significantly lower just prior to and for around 5 seconds after the response is made. **Figure 4H** shows mean aIC activity during the 3 second period after the choice was made (outcome period).

Within-subjects ANOVA using the factors Session (Late, Pun) and Choice (Alcohol, Social) revealed main effects of Choice (F(1,18)=10.7, p=0.004) and Session (F(1,18)=10.5, p=0.005), but no Choice by Session interaction (F(1,19)<1, p>0.05). Post-hoc t-tests show that aIC activity during the outcome period for Alcohol in Late choice is not different to Alcohol+shock in Punished Alcohol choice sessions (Late Alcohol mean 5.25 ± 2.8, Punished Alcohol mean 4.23 ± 3.29; paired t-test Late vs.

Pun: t(18)=1.32, p>0.05), in contrast aIC activity during the outcome period for Social in Late choice is significantly higher than during Punished alcohol choice (Late choice mean 3.79 ± 2.72, Punished Alcohol mean 2.37 ± 1.19; paired t-test Late vs. Pun: t(20)=2.92, p=0.009).

### Sex differences in aIC activity

We performed a secondary analysis on aIC activity centred around trial start separating the data into Male or Female subjects (Fig. S3). These data show significant differences in the recorded aIC activity during the Late choice sessions. Specifically, we found that aIC activity during the cue period was significantly higher in female rats than male rats for trials where alcohol was chosen (Fig. S3C), and where social is chosen (Fig. S3D).

### Cognitive modelling of choice between alcohol and social reward

Initial analysis of raw response latencies (time from lever insertion to choice) revealed overall patterns of decision-making speed across conditions. While these latencies provide a general measure of decision efficiency (Choi et al., 2022), we subsequently employed LBA modelling to decompose these reaction times into specific cognitive components. Latency distribution for each session is shown in **Figures 5A – 5C**, for each condition there was a log-normal distribution (Early Alcohol R^2^ = 0.86; Late Alcohol R^2^ = 0.98; Punished Alcohol R^2^=0.72; Early Social R^2^=0.87; Late Social R^2^=0.93; Punish Social R^2^=0.95). Mean latency per rat is shown in **Figure 5D**. There was no difference in response latency in Early choice (Alcohol median 11.2 ± 10.4, Social median 11.1 ± 10.6; paired t-test Alcohol vs. Social: t(21)=-0.4, p>0.05), but Alcohol latency was significantly faster than Social in Late Choice (Alcohol median 3.0 ± 4.5, Social median 5.9 ± 7.1; paired t-test Alcohol vs. Social: t(20)=-3.0, p=0.007). In Punished alcohol choice, there was no difference in response latency between Alcohol and Social (Alcohol median 18.3 ± 19.9, Social median 14.9 ± 17.2; paired t-test Alcohol vs. Social: t(20)<1, p>0.05).

**Figure 5.**
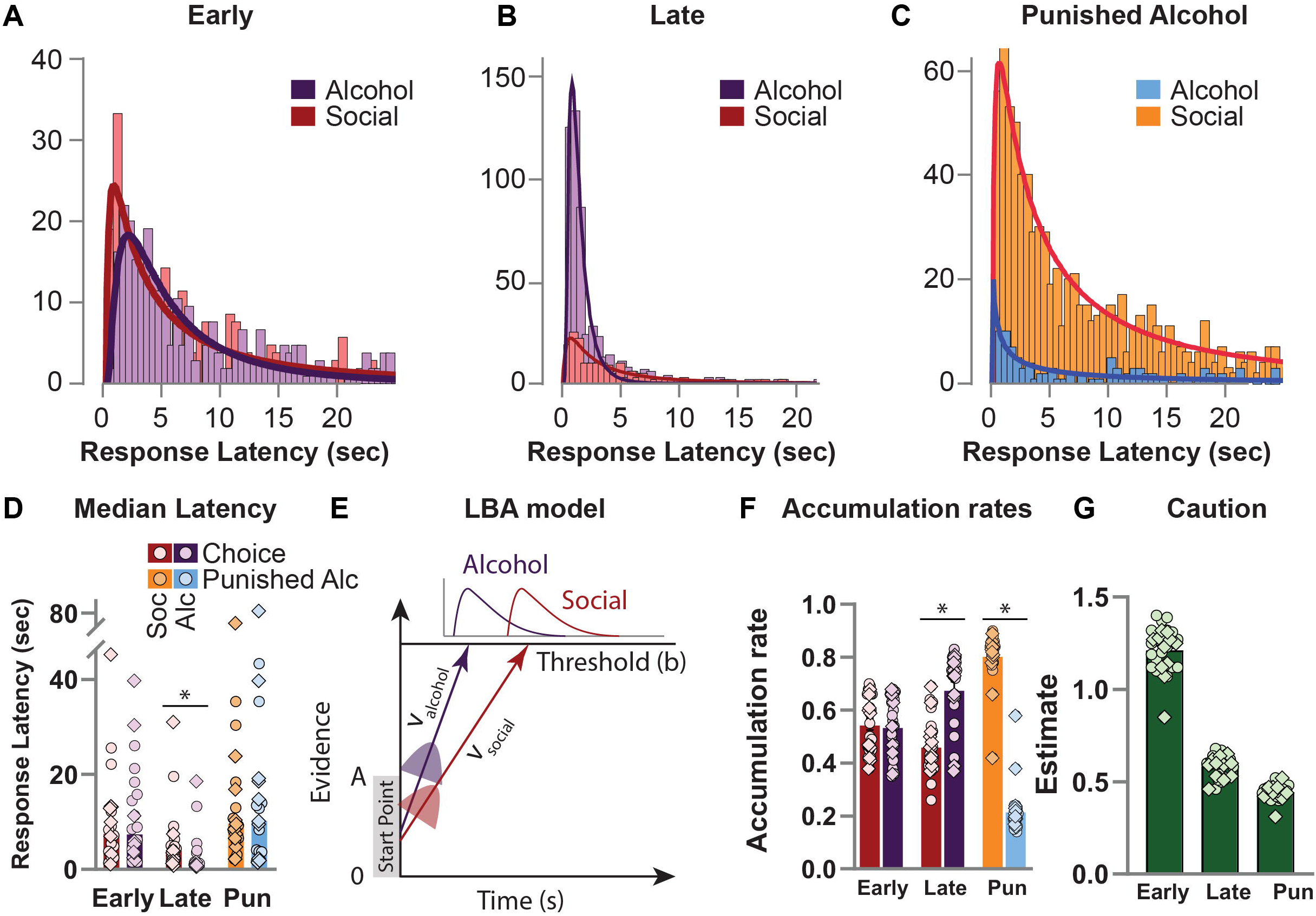
Cognitive modelling of choice behaviour. (**A, B, C**) Frequency distributions of response latency (from lever insertion) for Alcohol and Social choices across the three phases of the experiment. (**D**) Median Response Latency for either alcohol or social during the three phases of the experiment. (**E**) Model depicting the LBA model of choice. Salience (V) of alcohol or social accumulate separately to an evidence threshold (b) and thus to action selection. (**I**) Mean and individual subject data for accumulation rates for alcohol and social choice across the experiment. (**F**) Mean and individual subject data response caution across the experiment. *p<0.05. n=11 males; diamond shapes, n=12 females, circle shapes.

To elucidate the cognitive mechanisms underlying the rats’ choices between alcohol and social reward across the experimental phases, we applied we applied a systematic ‘power set’ of eight hierarchical Linear Ballistic Accumulator (LBA) models to the data (see Methods). We performed a formal model comparison and found that model M_SC, in which Stimulus Bias (accumulation rates) and Caution were allowed to vary across sessions, provided the best and most parsimonious account of the data (see Table 4 and Fig. S5B for comparison details). All subsequent results are based on this winning model. Model convergence was successful for all datasets, as indicated by R-hat (R^) values consistently below 1.05 (mean R^ = 1.000], range = [e.g., 1.000 - 1.000]) and effective sample sizes (ESS) well above conventional thresholds (mean ESS = 10340, min = 1489) for all key parameters (see example trace plots in **Figure S4**). However, posterior predictive checks revealed that while the model could successfully account for choice proportions (**Fig. S6A**), its ability to account for reaction times was limited, succeeding only for the stable, high-preference choices in the late acquisition phase (**Fig. S6B,C**). The model’s failure to capture reaction times in the punished condition, despite being the best-fitting model, suggests that punishment induces a qualitative shift in cognitive strategy that violates the core assumptions of the standard LBA framework.

### LBA Model Parameters Across Experimental Phases

The LBA model (**Figure 5E**) allowed us to extract estimates for key decision parameters: accumulation rates for alcohol (v*_alcohol_*) and social (v*_social_*) choices, start-point variability (A), threshold (b), and non-decision time (τ). **Figure 5F** shows the accumulation rate (v) for alcohol or social choices during the three phases. Based on the posterior distributions from the hierarchical model, we found no difference between the rates in early choice sessions (v*_alcohol_* mean 0.537 ± 0.109, v*_social_* mean 0.547 ± 0.107; paired t-test Alcohol vs. Social: t(22)=-0.218,p=0.8293 but in the late choice sessions v*_alcohol_* was significantly higher than v*_social_* (v*_alcohol_* mean 0.678 ± 0.140 v*_social_* mean 0.463 ± 0.106; paired t-test Alcohol vs. Social: t(22)=4.296,p<0.001). During the punished-alcohol sessions, v*_social_* was significantly higher than v*_alcohol_* (v*_alcohol_* mean 0.218 ± 0.092 v*_social_* mean 0.807 ± 0.099; paired t-test Alcohol vs. Social: t(22)=-14.8,p<0.001). This demonstrates that the model’s accumulation rate parameter successfully tracks the observed shifts in choice preference.

In stark contrast to the behavioural slowing observed in the punishment phase, the model revealed a paradoxical effect on decision caution. Caution is shown in **Figure 5G** and was found to significantly decrease from the Early choice sessions to the Late choice sessions (Early mean 0.607 ± 0.063, Late mean 0.295 ± 0.029; paired t-test Early vs Late: t(22)=21.5,p<0.001), and then decreased further in the Punished sessions (Punish mean 0.223 ± 0.020; paired t-test Late vs. Pun: t(22)=10.2,p<0.001). This counterintuitive result, where the model requires *less* caution to best explain behaviour that is demonstrably *slower* and more considered, indicates that the animals’ adaptation to punishment is not a simple modulation of a standard decision parameter, but a more complex shift in cognitive state that the model cannot capture.

### Relationship Between Choice Performance and Model-Derived Decision Bias

To quantify decision bias, we used a within-subjects estimation of the relative strength of evidence accumulation towards alcohol versus social reward (accumulation rate difference: v_alcohol –_ v_social_ ; hereafter Decision Bias). To provide a comparable metric in recorded aIC activity, for each rat we generated an ‘aIC difference’ score (aIC*_alcohol_* – aIC*_social_*). This measure reflects a within-subjects description of the relative magnitude of aIC activity during the cue period towards alcohol or social choices.

**Figure 6A** shows the Preference Score for the three phases of the experiment. We found a significant increase in preference score from Early to Late (Early mean –0.03 ± 0.53, Late mean 0.39 ± 0.57; paired t-test Early vs. Late: t(22)=-3.36, p=0.003), and a significant decrease in preference from Late to Pun (Pun mean -0.77 ± 0.36; paired t-test Late vs. Pun: t(22)=9.4, p<0.001). **Figure 6B** shows Decision Bias across the experimental phases. There was a significant increase in the Decision Bias towards positive values during the Late choice sessions (Early mean –0.01 ± 0.21, Late mean 0.21 ± 0.24; paired t-test Early vs. Late: t(22)=-4.4,p<0.001), indicating a faster evidence accumulation for alcohol choices in the Late choice sessions. In the punished-alcohol sessions, Decision Bias was negative, and significantly lower than Late Choice (Punish mean –0.589 ± 0.19; paired t-test Late vs. Pun: t(22)=14.7,p<0.001). **Figure 6C** shows the aIC Difference Score across the three phases. There is a significant difference between Early and Late choice sessions (Early mean 0.026 ± 1.13, Late mean 1.30 ± 1.57; paired t-test Early vs. Late: t(22)=-3.5, p=0.002) and Late choice and punished-alcohol sessions (Punish mean 0.21 ± 1.25; paired t-test Late vs. Pun: t(22)=3.28, p=0.003) but no difference between Early and Punished (paired t-test Early vs. Pun: t(22)=-0.5, p>0.05). This metric reveals a bias in aIC activity during the cue period for alcohol choices in the Late choice sessions. However, these findings also show that aIC activity does not increase for social choices during the Punished alcohol choice sessions.

**Figure 6.**
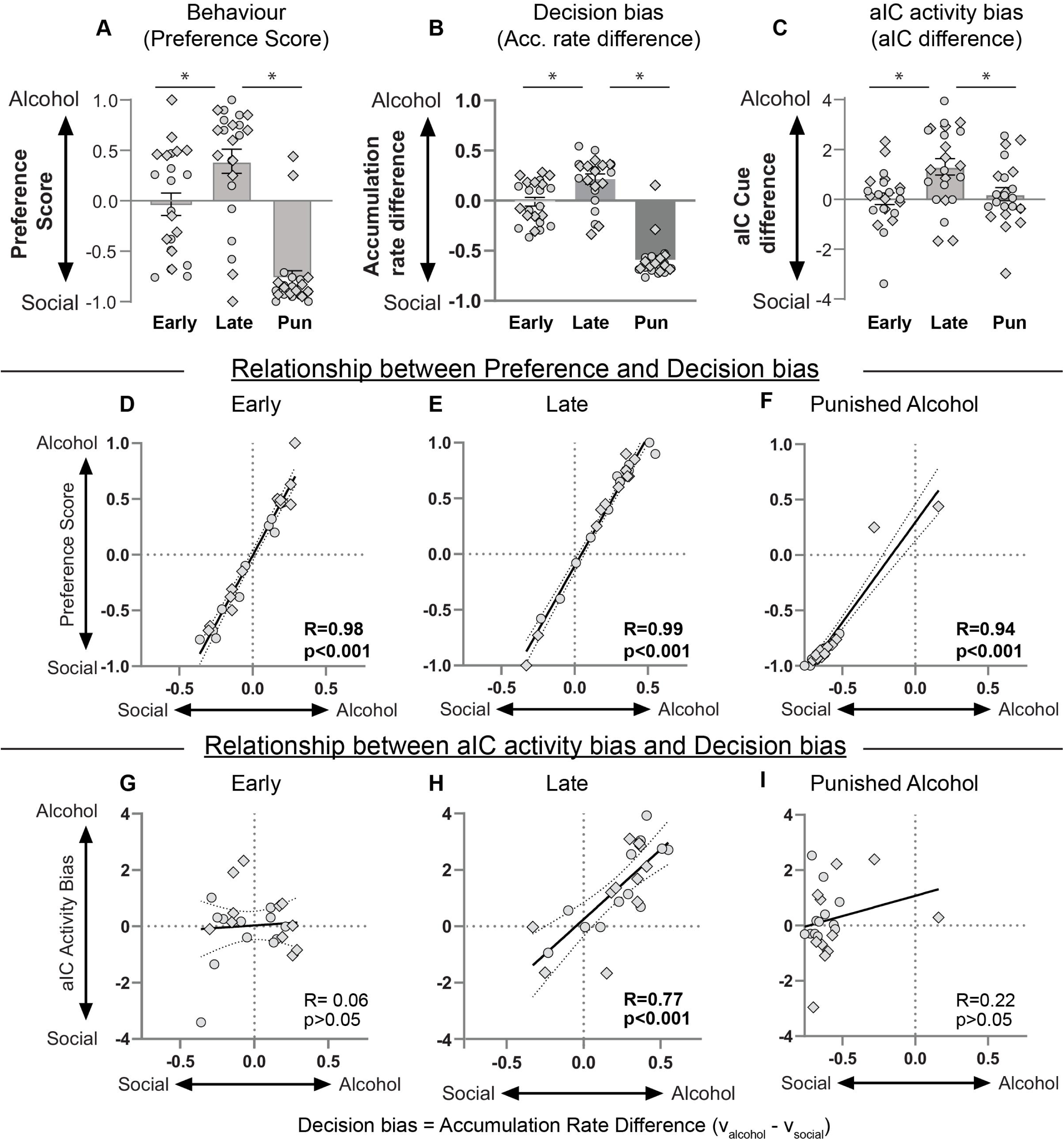
Relationship between decision-making and aIC calcium dynamics. (**A**) Behaviour: Preference score is a within-subjects value depicting the proportion of alcohol or social reward chosen in each phase of the experiment. (**B**) Decision Bias: Within-subjects value depicting the difference in accumulation rates for alcohol versus social (v_alcohol_ – v_social_). (**C**) aIC activity bias: Within-subject value depicting the difference in aIC activity during the cue period for trials where alcohol is chosen versus social (aIC_alcohol_ – aIC_social_). (**D, E, F**) Relationship between Preference and Decision bias. Correlations between Preference Score and Decision Bias for Early (**D**), Late (**E**), and Punished alcohol (**F**) sessions. (**G, H, I**) Relationship between aIC Activity Bias and Decision bias. Correlations between aIC and Decision Bias for Early (**G**), Late (**G**), and Punished alcohol (**I**) sessions. *p<0.05. n=11 males; diamond shapes, n=12 females, circle shapes.

We next examined the relationship between the behavioural observation, Preference Score, and the LBA-derived Decision Bias. We found a strong positive correlation between Preference Score and Decision Bias across each of the three phases shown in **Figure 6D-F**: Early: r=0.979,p<0.001; Late: r=0.987,p<0.001; Pun: r=0.950,p<0.001. These findings validate the Decision Bias parameter as a reliable metric capturing decision-making parameters underlying the expressed choice preferences.

### Relationship between Decision Bias and aIC activity during the cue period

Finally, we investigated the extent to which aIC calcium dynamics map onto these decision processes. We found that aIC Activity Bias was correlated with Decision Bias only in the Late choice sessions (**Figure 6H**, r=0.766,p<0.001). In contrast, this relationship was not significant in the Early choice sessions (**Figure 6G**, r=0.065,p=0.767) or in the Punished alcohol choice session (**Figure 6I**, r=0.223,p=0.307). In the model that we used for analysis, both accumulation rate and response caution varied between the experimental phases. As such, we conducted additional analysis testing whether response caution and Activity Bias were correlated in any phase. These results show that Activity Bias was not correlated with response caution in either Early (r=-0.379,p=0.074), Late (r=-0.259,p=0.233), or Punished alcohol (r=0.207,p=0.344) choice sessions, providing evidence that aIC activity during the cue period is related to faster choices for alcohol through a process of faster accumulation rate, rather than reduced decision threshold.

Correlations between aIC activity and individual accumulation rates or latencies, are shown in **Figure S7**. These data show that aIC calcium during the cue is significantly correlated with mean accumulation rate only for alcohol choices in the Late choice phase (**Fig. S7B**), and no other experimental phase, and not for social choices. These results provide further evidence for the relationship between alcohol preference and aIC activity during decision making.

To test whether aIC activity in the 10-sec period leading up to the choice response is better related to the accumulation rate than activity during the cue period, we also correlated these values (**Figure S7G-L**). These data show no relationship between these measures, indicating that aIC activity during the cue period is a better predictor of accumulation rate towards alcohol choices, than an equivalent period of aIC activity preceding the actual choice response. This dissociation provides strong evidence that aIC activity during the cue sets a ’Decision Bias’ that influences the subsequent evidence accumulation process, rather than tracking the accumulation as it unfolds.

## Discussion

In this study we measured aIC activity using calcium imaging with fiber photometry during choice between alcohol and social reward in a discrete-choice model (Lenoir et al., 2007). Replicating our (Marchant et al., 2023) and others (Augier et al., 2023) previous work, we show that both male and female rats show a choice preference for alcohol over social reward. We report four main findings. First, we show that aIC activity is higher for alcohol compared to social reward during both self-administration and choice. Second, emergence of an alcohol choice preference is associated with increased aIC activity during the cue period for trials where alcohol is chosen. Third, we show that punishment of the alcohol-reinforced response results in choice for social reward, which decreased cue-period aIC activity for alcohol choices, but had no change for social choices. Finally, cognitive modelling allowed us to describe the latent cognitive mechanisms underlying alcohol over social choices. Correlating the model-derived bias with activity revealed that aIC cue-period activity is related to Decision Bias for alcohol choices when preference is well-established. Overall, these findings highlight the specific role of aIC in the decision making in choice for alcohol.

### Consistent higher aIC activity towards alcohol reward compared to social reward

We found that aIC activity is more strongly associated with alcohol than social reward. We show that response-contingent alcohol elicits greater aIC calcium than social reward in both self-administration and choice. Anterior insula functions include interoception (Craig, 2009), emotional processing (Gu et al., 2013; Zych and Gogolla, 2021), and attentional modulation (Menon and Uddin, 2010; Zych and Gogolla, 2021). Greater aIC activity for alcohol compared to social outcomes may be a function of taste and visceral sensations. Insula cortex encodes visceral sensations, as well as both taste sensation and the stimuli which predict these outcomes (Samuelsen et al., 2012; Gardner and Fontanini, 2014). These observations typically come from IC regions more posterior than in this study, as visceral information from the body arrives in the posterior insula cortex (Allen et al., 1991). The rodent pIC typically shows higher activity for aversive tastes (Gehrlach et al., 2019). While aIC is also involved in aversive learning related to taste (Kayyal et al., 2019), neuroimaging studies show in humans that aIC is responsive to taste stimuli (Small, 2010; Rolls, 2016). One additional consideration is that social interaction involving touch also activates IC (Morrison et al., 2011; Gordon et al., 2013; Suvilehto et al., 2021), although again typically posterior IC. Regardless, one potential interpretational issue related to the observed aIC signal during social reward self-administration and choice is that the barrier prevents substantial touch, which may limit IC activation.

### Role of aIC in decision-making and choice

Several lines of research have identified a role for aIC in decision-making (Naqvi and Bechara, 2010; Daniel et al., 2017). Here we describe how decision processes change across the experimental phases. The LBA model specifically conceptualizes choice as a race between multiple independent accumulators which each gather evidence for a specific response (Forstmann et al., 2008; Donkin et al., 2011b). A key finding of our study is the correlation between aIC activity during the cue period and Decision Bias. This presents an important question regarding the temporal alignment of these events, as our model measures response latency from lever insertion, while the decision process likely begins at cue onset. We propose that cue-evoked aIC acts as a causal antecedent to the evidence accumulation process. This activity sets the initial parameters and motivational state governing subsequent race-to-threshold, effectively establishing decision bias before choice is executed. The importance of the cue period activity to this process is further exemplified by the observation that aIC activity in the 10 seconds preceding the choice action (i.e. lever press) does not correlate with the accumulation rate for any choice type.

The emergence of alcohol preference was characterized by a significant increase in decision bias towards alcohol. Conversely, punishment reversed Decision Bias, but also revealed a critical limitation of the model, for which the best fit required a paradoxical decrease in response caution, a finding at odds with observation. This likely reflects a fundamental shift in decision-making strategy, which may involve a switch to a different cognitive state not well-captured by the standard race model, which assumes a single cognitive strategy. One possibility is that, especially in punishment, the rats alternate between a task-engaged state and a hesitant state, resulting in a mix of fast and slow reaction times, as observed.

In the Late phase, a strong and stable alcohol preference may be representative of automatic choices where the preferred outcome reduces the need for prolonged deliberation before reaching a decision threshold (Donkin et al., 2011a; Shadlen and Shohamy, 2016; Redish et al., 2022). This form of decision-making would be associated with faster response times, which is observed for both alcohol and social choices. The observation that aIC activity tracks Decision Bias for alcohol, aligns with its proposed role in representing the motivational salience or expected value of outcomes (Menon and Uddin, 2010; Parkes and Balleine, 2013). In the Late phase, heightened and biased aIC activity might reflect an amplified salience attributed to alcohol cues, directly translating into a faster evidence accumulation for an alcohol choice. However, in the punished phase, the lack of correlation between aIC activity and Decision Bias, combined with the model’s failure to explain the extended latency with respect to increased caution, suggests that a different decision-making process is employed, which is not related to aIC activity. Consistent with this, aIC activity during the cue period is decoupled from Decision Bias. While we did not measure freezing in this experiment, previous work has shown that punishment is unlike fear conditioning and produces very little freezing particularly after learning (Bolles et al., 1980; Jean-Richard Dit-Bressel and McNally, 2015; Jean-Richard-Dit-Bressel et al., 2018). An additional argument against freezing is that behavioural responses were reallocated towards the social lever by the fourth session. The modelling data indicates that aIC is more related to the expression of preference-driven responses rather than the decision-making processes engaged when the alcohol choice is punished. Some studies have proposed that IC activity is related to decision-making during conflict (Naqvi et al., 2014; Daniel et al., 2017). However, these data show that aIC activity was more closely related to faster choices after preference is established.

The role of choice in addiction is not without controversy. The extent to which individuals with AUD seemingly maintain the ability to voluntarily continue alcohol use has been used to argue against the brain disease model of addiction (Heyman, 2021). However, the presence of choice does not exclude the possibility that the choices are influenced by the neurobiological consequences of alcohol use (Heilig et al., 2021; Pickard, 2022), and there are many plausible dysfunctions in decision-making in AUD (Verdejo-Garcia et al., 2018). Value-based decision making can be a valuable framework to describe a pathway towards recovery from substance use disorders (Field et al., 2020), and potential decision processes which result in persistent alcohol use (Redish et al., 2008). The use of LBA modelling in this study aligns with the pursuit of computational validity (Redish et al., 2022), which emphasizes that translating behavioural findings across different contexts or species should be grounded in similarity of the underlying information processing and computational algorithms, rather than task resemblances or general theoretical constructs. By decomposing behaviour into quantifiable model-based parameters such as decision bias and response caution, we move beyond a descriptive account of choice to probe the latent computational mechanisms driving those choices (Donkin et al., 2011b). It will be of interest in future studies to determine whether comparable metrics are observed in human laboratory tasks (e.g. (Li et al., 2020; Karlsson et al., 2025)).

Previous work has shown that activity in the rat IC, more posterior to recorded here, responds both to consumed rewards as well as cues which predict their arrival (Samuelsen et al., 2012; Gardner and Fontanini, 2014). This study expands on these observations to show that activity in anterior IC tracks cue-induced response anticipation, and decision-making during choice. Activity in aIC contributes towards choice between goal-directed actions based on aIC-dependent representation of value (Parkes and Balleine, 2013; Parkes et al., 2015). Communication between aIC and the nucleus accumbens (NAc) core is critical for retrieving outcome values and directing choice (Parkes et al., 2015). Inhibition of the aIC to NAc pathway decreases aversion-resistant alcohol use (Seif et al., 2013), and chemogenetic stimulation increases alcohol consumption (Haaranen et al., 2020). In rodents, alcohol dependence is associated with altered function connectivity between IC and NAc (Scuppa et al., 2020), and increased connectivity between aIC and NAc is associated with AUD in humans (Grodin et al., 2018; Le et al., 2022).

### Concluding remarks

In this study we show how activity in aIC relates to choice between alcohol and social reward. We found that aIC activity is consistently higher for alcohol reward both during self-administration and choice. Our modelling of the decision-making process showed that differential activity during the decision period, particularly when preference is established, reflects a stronger evidence accumulation bias towards alcohol.

## Acknowledgments

The authors gratefully acknowledge: Dr. Philip Jean-Richard-Dit-Bressel for assistance with photometry analysis and sharing early versions of the scripts we used; Professor Andrew Heathcote for advice on the modelling approach; the VUmc Histology Imaging Unit for their support & assistance in whole-slide imaging.

## Author’s contributions

NJM & TDV conceived of the experiments; YvM, DS, IB, MES conducted the experiments; NJM wrote the first draft of the manuscript; All authors gave final approval of the submission.

## References

Allen GV, Saper CB, Hurley KM, Cechetto DF (1991) Organization of visceral and limbic connections in the insular cortex of the rat. J Comp Neurol 311:1–16.

Annis J, Miller BJ, Palmeri TJ (2017) Bayesian inference with Stan: A tutorial on adding custom distributions. Behav Res Methods 49:863–886.

Augier E, Barbier E, Dulman RS, Licheri V, Augier G, Domi E, Barchiesi R, Farris S, Natt D, Mayfield RD, Adermark L, Heilig M (2018) A molecular mechanism for choosing alcohol over an alternative reward. Science 360:1321–1326.

Augier G, Schwabl V, Lguensat A, Atudorei M, Iyere OC, Solander SE, Augier E (2023) Wistar rats choose alcohol over social interaction in a discrete-choice model. Neuropsychopharmacology 48:1098–1107.

Bankhead P, Loughrey MB, Fernandez JA, Dombrowski Y, McArt DG, Dunne PD, McQuaid S, Gray RT, Murray LJ, Coleman HG, James JA, Salto-Tellez M, Hamilton PW (2017) QuPath: Open source software for digital pathology image analysis. Sci Rep 7:16878.

Belin-Rauscent A, Daniel ML, Puaud M, Jupp B, Sawiak S, Howett D, McKenzie C, Caprioli D, Besson M, Robbins TW, Everitt BJ, Dalley JW, Belin D (2016) From impulses to maladaptive actions: the insula is a neurobiological gate for the development of compulsive behavior. Mol Psychiatry 21:491–499.

Bickel WK, Johnson MW, Koffarnus MN, MacKillop J, Murphy JG (2014) The behavioral economics of substance use disorders: reinforcement pathologies and their repair. Annu Rev Clin Psychol 10:641–677.

Bolles RC, Holtz R, Dunn T, Hill W (1980) Comparisons of stimulus learning and response learning in a punishment situation. Learning and Motivation 11:78–96.

Brown SD, Heathcote A (2008) The simplest complete model of choice response time: linear ballistic accumulation. Cogn Psychol 57:153–178.

Campbell EJ, Flanagan JPM, Walker LC, Hill M, Marchant NJ, Lawrence AJ (2019) Anterior Insular Cortex is Critical for the Propensity to Relapse Following Punishment-Imposed Abstinence of Alcohol Seeking. J Neurosci 39:1077–1087.

Cantin L, Lenoir M, Augier E, Vanhille N, Dubreucq S, Serre F, Vouillac C, Ahmed SH (2010) Cocaine is low on the value ladder of rats: possible evidence for resilience to addiction. PLoS One 5:e11592.

Caprioli D, Venniro M, Zeric T, Li X, Adhikary S, Madangopal R, Marchant NJ, Lucantonio F, Schoenbaum G, Bossert JM, Shaham Y (2015) Effect of the novel positive allosteric modulator of mGluR 2 AZD8529 on incubation of methamphetamine craving after prolonged voluntary abstinence in a rat model. Biological Psychiatry.

Choi EA, Husic M, Millan EZ, Gilchrist S, Power JM, Jean-Richard Dit Bressel P, McNally GP (2022) A Corticothalamic Circuit Trades off Speed for Safety during Decision-Making under Motivational Conflict. J Neurosci 42:3473–3483.

Chow JJ, Beacher NJ, Chabot JM, Oke M, Venniro M, Lin DT, Shaham Y (2022) Characterization of operant social interaction in rats: effects of access duration, effort, peer familiarity, housing conditions, and choice between social interaction vs. food or remifentanil. Psychopharmacology (Berl) 239:2093–2108.

Craig AD (2009) How do you feel - now? The anterior insula and human awareness. Nature Reviews Neuroscience 10:59–70.

Daniel ML, Cocker PJ, Lacoste J, Mar AC, Houeto JL, Belin-Rauscent A, Belin D (2017) The anterior insula bidirectionally modulates cost-benefit decision-making on a rodent gambling task. Eur J Neurosci 46:2620–2628.

Domi E, Xu L, Toivainen S, Nordeman A, Gobbo F, Venniro M, Shaham Y, Messing RO, Visser E, van den Oever MC, Holm L, Barbier E, Augier E, Heilig M (2021) A neural substrate of compulsive alcohol use. Sci Adv 7.

Donkin C, Brown S, Heathcote A (2011a) Drawing conclusions from choice response time models: A tutorial using the linear ballistic accumulator. Journal of Mathematical Psychology 55:140–151.

Donkin C, Brown S, Heathcote A, Wagenmakers E-J (2011b) Diffusion versus linear ballistic accumulation: different models but the same conclusions about psychological processes? Psychonomic Bulletin & Review 18:61–69.

Field M, Heather N, Murphy JG, Stafford T, Tucker JA, Witkiewitz K (2020) Recovery from addiction: Behavioral economics and value-based decision making. Psychol Addict Behav 34:182–193.

Forget B, Pushparaj A, Le Foll B (2010) Granular insular cortex inactivation as a novel therapeutic strategy for nicotine addiction. Biol Psychiatry 68:265–271.

Forstmann BU, Dutilh G, Brown S, Neumann J, von Cramon DY, Ridderinkhof KR, Wagenmakers EJ (2008) Striatum and pre-SMA facilitate decision-making under time pressure. Proc Natl Acad Sci U S A 105:17538–17542.

Gardner MP, Fontanini A (2014) Encoding and tracking of outcome-specific expectancy in the gustatory cortex of alert rats. J Neurosci 34:13000–13017.

Gehrlach DA, Dolensek N, Klein AS, Roy Chowdhury R, Matthys A, Junghanel M, Gaitanos TN, Podgornik A, Black TD, Reddy Vaka N, Conzelmann KK, Gogolla N (2019) Aversive state processing in the posterior insular cortex. Nat Neurosci 22:1424–1437.

Gelman A, Rubin DB (1992) Inference from Iterative Simulation Using Multiple Sequences. Statistical Science 7:457–472, 416.

Gelman A, Carlin JB, Stern HS, Dunson DB, Vehtari A, Rubin DB (2013) Bayesian Data Analysis (3rd ed.). CRC Press.

Ghareh H, Alonso-Lozares I, Schetters D, Herman RJ, Heistek TS, Van Mourik Y, Jean-Richard-Dit-Bressel P, Zernig G, Mansvelder HD, De Vries TJ, Marchant NJ (2022) Role of anterior insula cortex in context-induced relapse of nicotine-seeking. Elife 11.

Gilpin NW, Koob GF (2008) Neurobiology of alcohol dependence: focus on motivational mechanisms. Alcohol Res Health 31:185–195.

Goldstein RZ, Volkow ND (2011) Dysfunction of the prefrontal cortex in addiction: neuroimaging findings and clinical implications. Nat Rev Neurosci 12:652–669.

Gordon I, Voos AC, Bennett RH, Bolling DZ, Pelphrey KA, Kaiser MD (2013) Brain mechanisms for processing affective touch. Hum Brain Mapp 34:914–922.

Grodin EN, Sussman L, Sundby K, Brennan GM, Diazgranados N, Heilig M, Momenan R (2018) Neural Correlates of Compulsive Alcohol Seeking in Heavy Drinkers. Biol Psychiatry Cogn Neurosci Neuroimaging 3:1022–1031.

Gu X, Hof PR, Friston KJ, Fan J (2013) Anterior insular cortex and emotional awareness. J Comp Neurol 521:3371–3388.

Haaranen M, Schafer A, Jarvi V, Hyytia P (2020) Chemogenetic Stimulation and Silencing of the Insula, Amygdala, Nucleus Accumbens, and Their Connections Differentially Modulate Alcohol Drinking in Rats. Front Behav Neurosci 14:580849.

Hall W, Carter A, Forlini C (2015) The brain disease model of addiction: is it supported by the evidence and has it delivered on its promises? Lancet Psychiatry 2:105–110.

Hart CL (2017) Viewing addiction as a brain disease promotes social injustice. Nat Hum Behav 1.

Heather N (1998) A conceptual framework for explaining drug addiction. J Psychopharmacol 12:3–7.

Heatherton TF, Wagner DD (2011) Cognitive neuroscience of self-regulation failure. Trends Cogn Sci 15:132–139.

Heilig M, Epstein DH, Nader MA, Shaham Y (2016) Time to connect: bringing social context into addiction neuroscience. Nat Rev Neurosci 17:592–599.

Heilig M, MacKillop J, Martinez D, Rehm J, Leggio L, Vanderschuren L (2021) Addiction as a brain disease revised: why it still matters, and the need for consilience. Neuropsychopharmacology 46:1715–1723.

Heilig M, Barbier E, Johnstone AL, Tapocik J, Meinhardt MW, Pfarr S, Wahlestedt C, Sommer WH (2017) Reprogramming of mPFC transcriptome and function in alcohol dependence. Genes Brain Behav 16:86–100.

Heyman GM (2021) How individuals make choices explains addiction’s distinctive, non-eliminable features. Behav Brain Res 397:112899.

Hoffman MD, Gelman A (2014) The No-U-Turn Sampler: Adaptively Setting Path Lengths in Hamiltonian Monte Carlo. J Mach Learn Res 15:1593–1623.

Hunt GM, Azrin NH (1973) A community-reinforcement approach to alcoholism. Behav Res Ther 11:91–104.

Janes AC, Krantz NL, Nickerson LD, Frederick BB, Lukas SE (2020) Craving and Cue Reactivity in Nicotine-Dependent Tobacco Smokers Is Associated With Different Insula Networks. Biol Psychiatry Cogn Neurosci Neuroimaging 5:76–83.

Janes AC, Gilman JM, Radoman M, Pachas G, Fava M, Evins AE (2017) Revisiting the role of the insula and smoking cue-reactivity in relapse: A replication and extension of neuroimaging findings. Drug Alcohol Depend 179:8–12.

Jean-Richard-Dit-Bressel P, Killcross S, McNally GP (2018) Behavioral and neurobiological mechanisms of punishment: implications for psychiatric disorders. Neuropsychopharmacology 43:1639–1650.

Jean-Richard-Dit-Bressel P, Clifford CWG, McNally GP (2020) Analyzing Event-Related Transients: Confidence Intervals, Permutation Tests, and Consecutive Thresholds. Front Mol Neurosci 13:14.

Jean-Richard Dit-Bressel P, McNally GP (2015) The role of the basolateral amygdala in punishment. Learning & Memory 22:128–137.

Jentsch JD, Taylor JR (1999) Impulsivity resulting from frontostriatal dysfunction in drug abuse: implications for the control of behavior by reward-related stimuli. Psychopharmacology (Berl) 146:373–390.

Jones JA, Belin-Rauscent A, Jupp B, Fouyssac M, Sawiak SJ, Zuhlsdorff K, Zhukovsky P, Hebdon L, Velazquez Sanchez C, Robbins TW, Everitt BJ, Belin D, Dalley JW (2024) Neurobehavioral Precursors of Compulsive Cocaine Seeking in Dual Frontostriatal Circuits. Biol Psychiatry Glob Open Sci 4:194–202.

Joshi DD, Puaud M, Fouyssac M, Belin-Rauscent A, Everitt B, Belin D (2020) The anterior insular cortex in the rat exerts an inhibitory influence over the loss of control of heroin intake and subsequent propensity to relapse. Eur J Neurosci 52:4115–4126.

Joutsa J, Moussawi K, Siddiqi SH, Abdolahi A, Drew W, Cohen AL, Ross TJ, Deshpande HU, Wang HZ, Bruss J, Stein EA, Volkow ND, Grafman JH, van Wijngaarden E, Boes AD, Fox MD (2022) Brain lesions disrupting addiction map to a common human brain circuit. Nat Med 28:1249–1255.

Karlsson H, McNtyre S, Gustavson S, Andersson D, Szczot I, Heilig M, Perini I (2025) Choice of alcohol over a natural reward: an experimental study in light and heavy social drinkers. Psychopharmacology (Berl) 242:327–336.

Kayyal H, Yiannakas A, Kolatt Chandran S, Khamaisy M, Sharma V, Rosenblum K (2019) Activity of Insula to Basolateral Amygdala Projecting Neurons is Necessary and Sufficient for Taste Valence Representation. J Neurosci 39:9369–9382.

Le AD, Kiianmaa K, Cunningham CL, Engel JA, Ericson M, Soderpalm B, Koob GF, Roberts AJ, Weiss F, Hyytia P, Janhunen S, Mikkola J, Backstrom P, Ponomarev I, Crabbe JC (2001) Neurobiological processes in alcohol addiction. Alcohol Clin Exp Res 25:144S–151S.

Le TM, Malone T, Li CR (2022) Positive alcohol expectancy and resting-state functional connectivity of the insula in problem drinking. Drug Alcohol Depend 231:109248.

Lenoir M, Serre F, Cantin L, Ahmed SH (2007) Intense sweetness surpasses cocaine reward. PLoS One 2:e698.

Leshner AI (1997) Addiction is a brain disease, and it matters. Science 278:45–47.

Li J, Murray CH, Weafer J, de Wit H (2020) Subjective Effects of Alcohol Predict Alcohol Choice in Social Drinkers. Alcohol Clin Exp Res 44:2579–2587.

Marchant NJ, Hamlin AS, McNally GP (2009) Lateral hypothalamus is required for context-induced reinstatement of extinguished reward seeking. J Neurosci 29:1331–1342.

Marchant NJ, Furlong TM, McNally GP (2010) Medial dorsal hypothalamus mediates the inhibition of reward seeking after extinction. J Neurosci 30:14102–14115.

Marchant NJ, Campbell EJ, Kaganovsky K (2018) Punishment of alcohol-reinforced responding in alcohol preferring P rats reveals a bimodal population: Implications for models of compulsive drug seeking. Prog Neuropsychopharmacol Biol Psychiatry 87:68–77.

Marchant NJ, McDonald AJ, Matsuzaki R, van Mourik Y, Schetters D, De Vries TJ (2023) Rats choose alcohol over social reward in an operant choice procedure. Neuropsychopharmacology 48:585–593.

McDonald AJ, Nemat P, van ’t Hullenaar T, Schetters D, van Mourik Y, Alonso-Lozares I, De Vries TJ, Marchant NJ (2024) Punishment-resistant alcohol intake is mediated by the nucleus accumbens shell in female rats. Neuropsychopharmacology 49:2022–2031.

McNally GP (2021) Motivational competition and the paraventricular thalamus. Neurosci Biobehav Rev 125:193–207.

Menon V, Uddin LQ (2010) Saliency, switching, attention and control: a network model of insula function. Brain Struct Funct 214:655–667.

Miller NE (1971) Neal E. Miller: Selected papers. Chicago, IL.: Aldine Atherton Inc.

Morrison I, Bjornsdotter M, Olausson H (2011) Vicarious responses to social touch in posterior insular cortex are tuned to pleasant caressing speeds. J Neurosci 31:9554–9562.

Naqvi NH, Bechara A (2010) The insula and drug addiction: an interoceptive view of pleasure, urges, and decision-making. Brain Struct Funct 214:435–450.

Naqvi NH, Rudrauf D, Damasio H, Bechara A (2007) Damage to the insula disrupts addiction to cigarette smoking. Science 315:531–534.

Naqvi NH, Gaznick N, Tranel D, Bechara A (2014) The insula: a critical neural substrate for craving and drug seeking under conflict and risk. Ann N Y Acad Sci 1316:53–70.

Parkes SL, Balleine BW (2013) Incentive memory: evidence the basolateral amygdala encodes and the insular cortex retrieves outcome values to guide choice between goal-directed actions. J Neurosci 33:8753–8763.

Parkes SL, Bradfield LA, Balleine BW (2015) Interaction of insular cortex and ventral striatum mediates the effect of incentive memory on choice between goal-directed actions. J Neurosci 35:6464–6471.

Paxinos G, Watson C (2008) The rat brain in stereotaxic coordinates. Sixth edition., 3 Edition. San Diego, CA: Academic Press.

Pickard H (2022) Is addiction a brain disease? A plea for agnosticism and heterogeneity. Psychopharmacology (Berl) 239:993–1007.

Pickard H, Ahmed SH, Foddy B (2015) Alternative models of addiction. Front Psychiatry 6:20.

Prendergast M, Podus D, Finney J, Greenwell L, Roll J (2006) Contingency management for treatment of substance use disorders: a meta-analysis. Addiction 101:1546–1560.

Pushparaj A, Hamani C, Yu W, Shin DS, Kang B, Nobrega JN, Le Foll B (2013) Electrical stimulation of the insular region attenuates nicotine-taking and nicotine-seeking behaviors. Neuropsychopharmacology 38:690–698.

Redish AD, Jensen S, Johnson A (2008) A unified framework for addiction: vulnerabilities in the decision process. Behav Brain Sci 31:415–437; discussion 437–487.

Redish AD, Kepecs A, Anderson LM, Calvin OL, Grissom NM, Haynos AF, Heilbronner SR, Herman AB, Jacob S, Ma S, Vilares I, Vinogradov S, Walters CJ, Widge AS, Zick JL, Zilverstand A (2022) Computational validity: using computation to translate behaviours across species. Philos Trans R Soc Lond B Biol Sci 377:20200525.

Rolls ET (2016) Functions of the anterior insula in taste, autonomic, and related functions. Brain Cogn 110:4–19.

Rotge JY, Cocker PJ, Daniel ML, Belin-Rauscent A, Everitt BJ, Belin D (2017) Bidirectional regulation over the development and expression of loss of control over cocaine intake by the anterior insula. Psychopharmacology (Berl) 234:1623–1631.

Salvatier J, Wiecki TV, Fonnesbeck C (2016) Probabilistic programming in Python using PyMC3. Peerj Comput Sci.

Samuelsen CL, Gardner MP, Fontanini A (2012) Effects of cue-triggered expectation on cortical processing of taste. Neuron 74:410–422.

Scuppa G, Tambalo S, Pfarr S, Sommer WH, Bifone A (2020) Aberrant insular cortex connectivity in abstinent alcohol-dependent rats is reversed by dopamine D3 receptor blockade. Addict Biol 25:e12744.

Seif T, Chang SJ, Simms JA, Gibb SL, Dadgar J, Chen BT, Harvey BK, Ron D, Messing RO, Bonci A, Hopf FW (2013) Cortical activation of accumbens hyperpolarization-active NMDARs mediates aversion-resistant alcohol intake. Nat Neurosci 16:1094–1100.

Shadlen MN, Shohamy D (2016) Decision Making and Sequential Sampling from Memory. Neuron 90:927–939.

Small DM (2010) Taste representation in the human insula. Brain Struct Funct 214:551–561.

Suvilehto JT, Renvall V, Nummenmaa L (2021) Relationship-specific Encoding of Social Touch in Somatosensory and Insular Cortices. Neuroscience 464:105–116.

Vandaele Y, Daeppen JB (2022) From concepts to treatment: a dialog between a preclinical researcher and a clinician in addiction medicine. Transl Psychiatry 12:401.

Venniro M, Russell TI, Zhang M, Shaham Y (2019) Operant Social Reward Decreases Incubation of Heroin Craving in Male and Female Rats. Biol Psychiatry 86:848–856.

Venniro M, Zhang M, Caprioli D, Hoots JK, Golden SA, Heins C, Morales M, Epstein DH, Shaham Y (2018) Volitional social interaction prevents drug addiction in rat models. Nat Neurosci 21:1520–1529.

Venniro M, Caprioli D, Zhang M, Whitaker LR, Zhang S, Warren BL, Cifani C, Marchant NJ, Yizhar O, Bossert JM, Chiamulera C, Morales M, Shaham Y (2017) The Anterior Insular Cortex-->Central Amygdala Glutamatergic Pathway Is Critical to Relapse after Contingency Management. Neuron 96:414–427 e418.

Verdejo-Garcia A, Chong TT, Stout JC, Yucel M, London ED (2018) Stages of dysfunctional decision-making in addiction. Pharmacol Biochem Behav 164:99–105.

Zych AD, Gogolla N (2021) Expressions of emotions across species. Curr Opin Neurobiol 68:57–66.

