## Supplementary figures and images for "Anterior insula activity during alcohol and social reward self-administration and choice in male and female rats"

### Figure S1

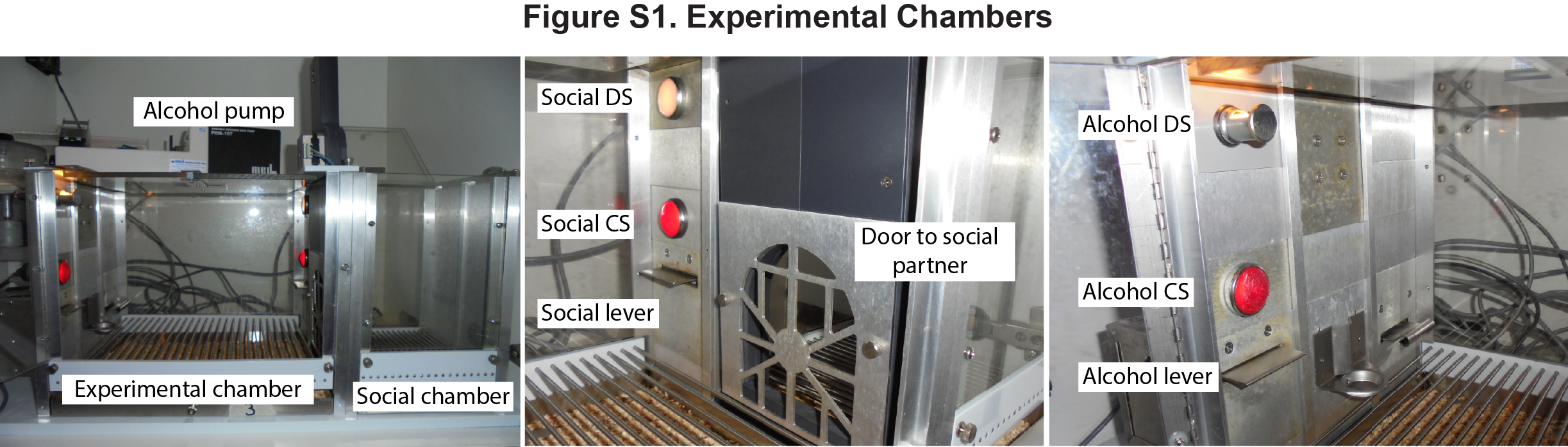

### Figure S2

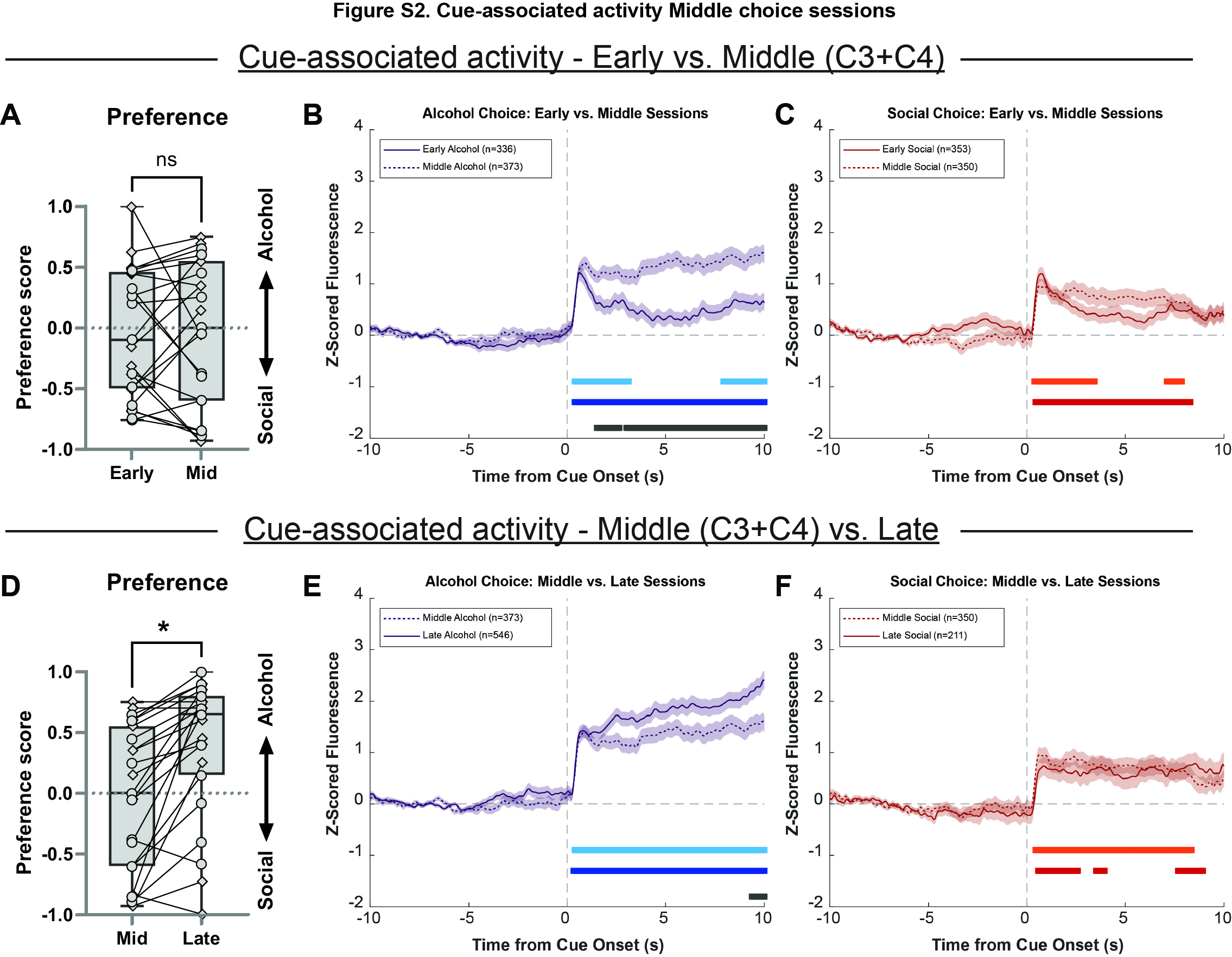

### Figure S3

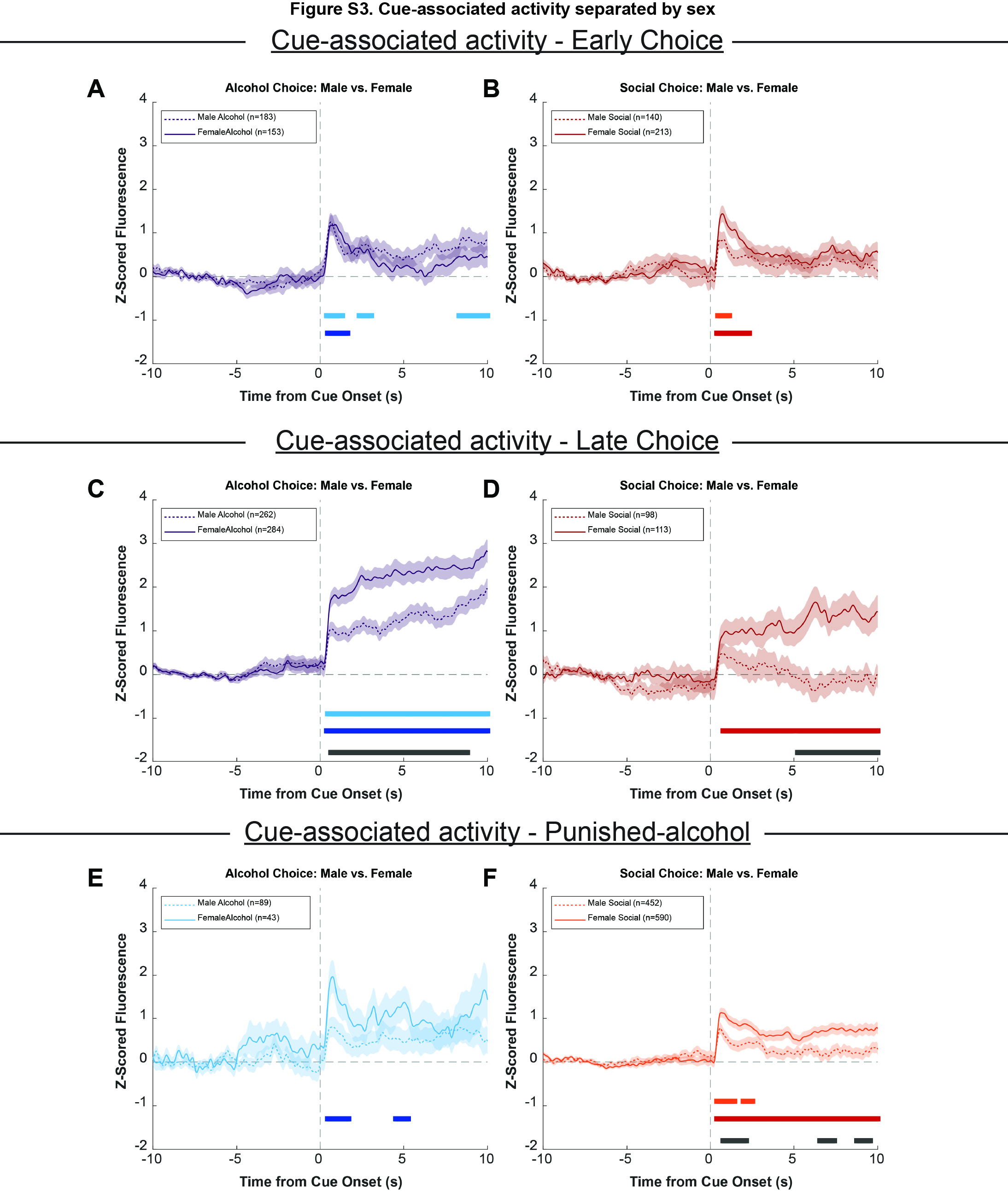

### Figure S4

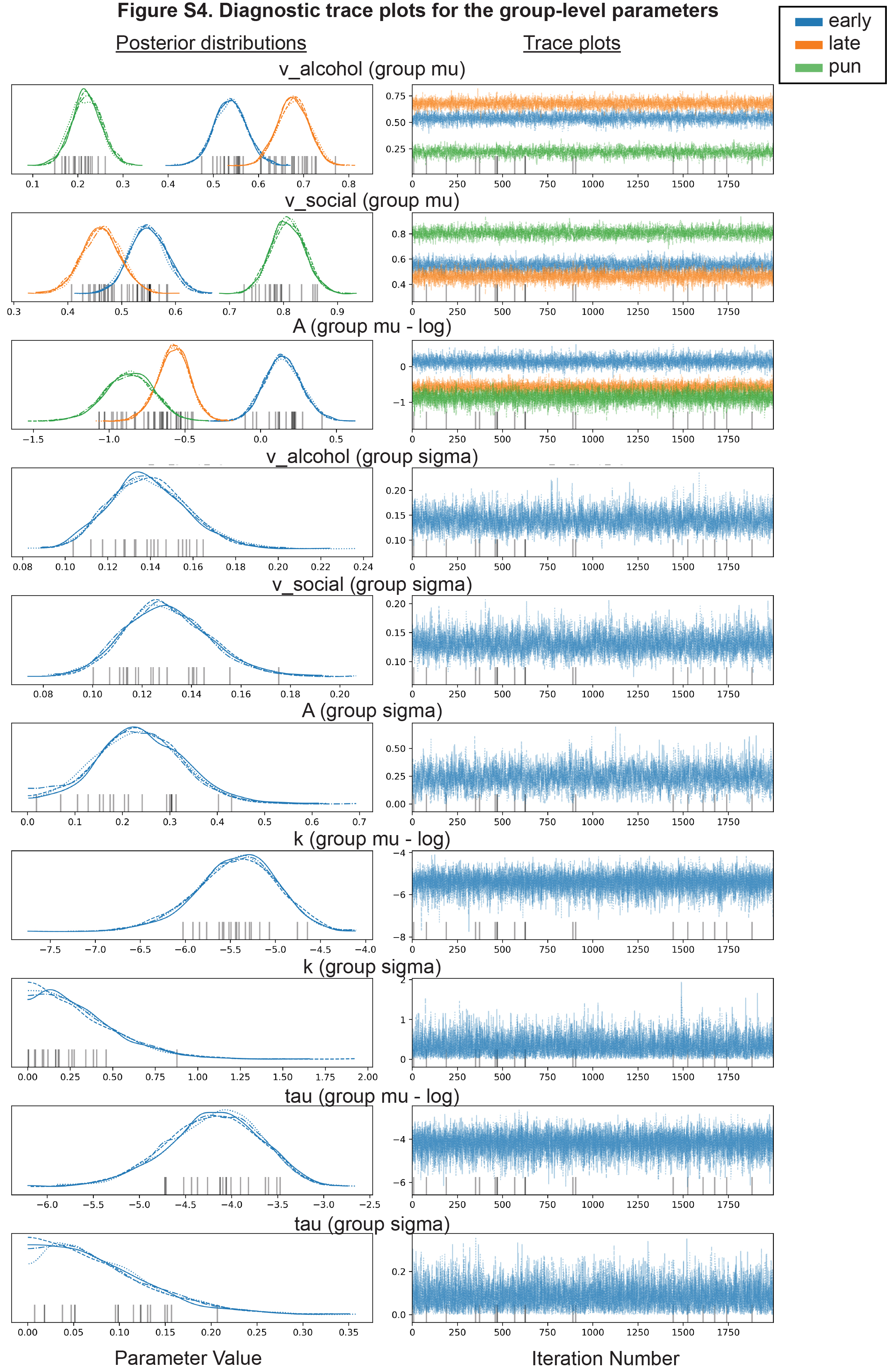

### Figure S5

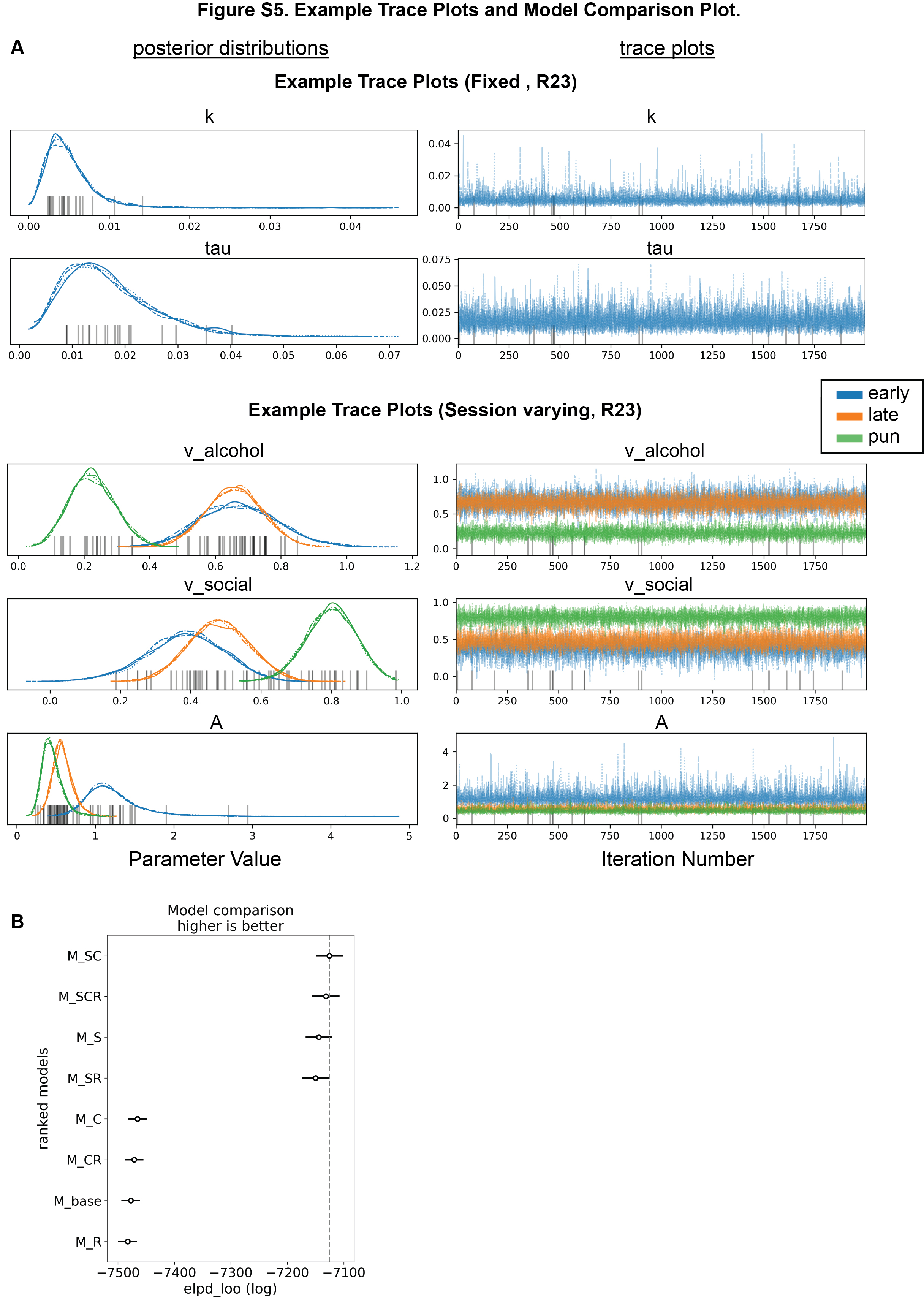

### Figure S6

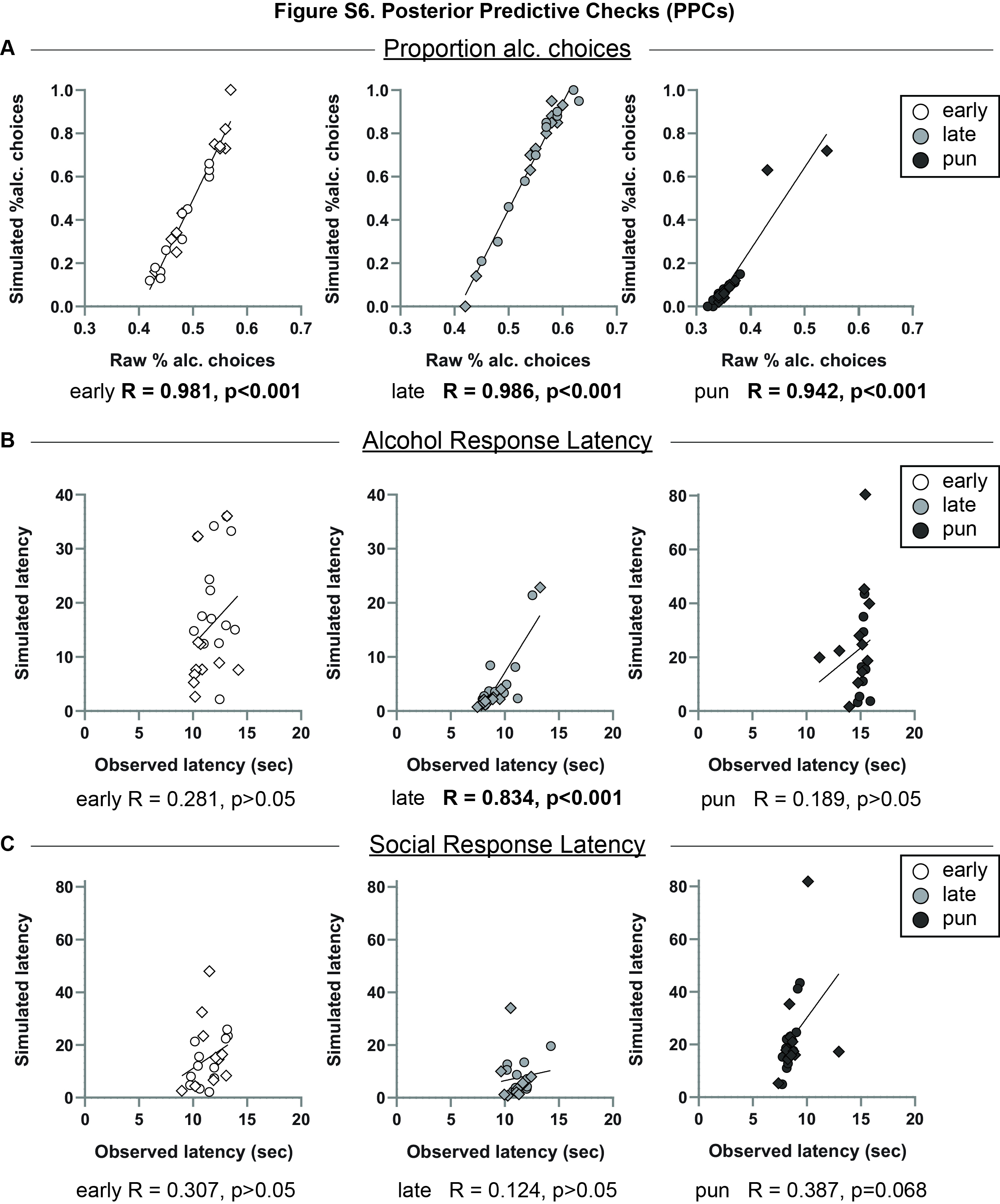

### Figure S7

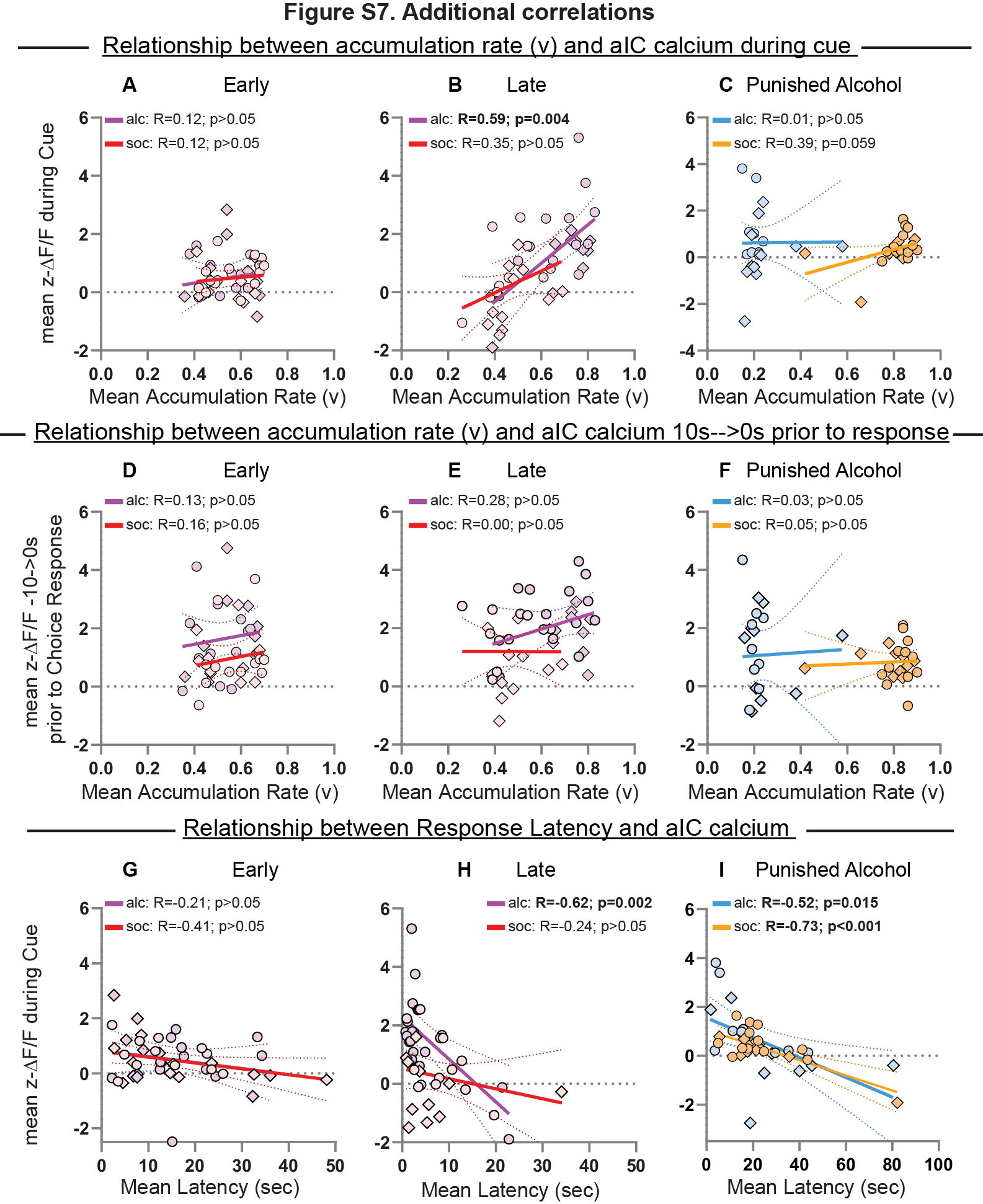
